# BMI1 regulates human erythroid self-renewal through both gene repression and gene activation

**DOI:** 10.1101/2024.02.02.578704

**Authors:** Kathleen E. McGrath, Anne D. Koniski, Kristin Murphy, Michael Getman, Hyun Hyung An, Vincent P. Schulz, Ah Ram Kim, Bin Zhang, Taylor L. Schofield, Julien Papoin, Lionel Blanc, Paul D. Kingsley, Connie M. Westhoff, Patrick G. Gallagher, Stella T. Chou, Laurie A. Steiner, James Palis

**Affiliations:** Department of Pediatrics, University of Rochester Medical Center, Rochester, NY USA; Dept. of Pediatrics, The Children’s Hospital of Philadelphia, Philadelphia, PA, USA; Dept. of Pediatrics, Yale School of Medicine, New Haven, CT, USA; Department of Pathology and Laboratory Medicine, University of Rochester Medical Center, Rochester, NY, USA; Institute of Molecular Medicine, Feinstein Institutes for Medical Research, Manhasset, NY, USA; Immunohematology and Genomics, New York Blood Center, New York, NY, USA; Nationwide Children’s Hospital, Ohio State University, Columbus, OH, USA

## Abstract

The limited proliferative capacity of erythroid precursors is a major obstacle to generate sufficient numbers of in vitro-derived red blood cells (RBC) for clinical purposes. We and others have determined that BMI1, a member of the polycomb repressive complex 1 (PRC1), is both necessary and sufficient to drive extensive proliferation of self-renewing erythroblasts (SREs). However, the mechanisms of BMI1 action remain poorly understood. BMI1 overexpression led to 10 billion-fold increase BMI1-induced (i)SRE self-renewal. Despite prolonged culture and BMI1 overexpression, human iSREs can terminally mature and agglutinate with typing reagent monoclonal antibodies against conventional RBC antigens. BMI1 and RING1B occupancy, along with repressive histone marks, were identified at known BMI1 target genes, including the INK-ARF locus, consistent with an altered cell cycle following BMI1 inhibition. We also identified upregulated BMI1 target genes with low repressive histone modifications, including key regulator of cholesterol homeostasis. Functional studies suggest that both cholesterol import and synthesis are essential for BMI1-associated self-renewal. These findings support the hypothesis that BMI1 regulates erythroid self-renewal not only through gene repression but also through gene activation and offer a strategy to expand the pool of immature erythroid precursors for eventual clinical uses.

## Introduction

Self-renewal, the generation of daughter cells with the same developmental potential as the parent cell, has been investigated within the hematopoietic system as a specific characteristic of hematopoietic stem cells (HSCs) (Bryder, 2006). However, hematopoietic progenitors and even some mature blood cells, such as subpopulations of tissue-resident macrophages and B1 lymphocytes, can also self-renew to maintain and expand their cellular compartments (Sieweke, 2013; Kobayashi, 2020). In the erythroid lineage, ex vivo culture of erythroid ‘progenitors’ leads to their limited, approximately 200-300-fold, expansion in the presence of erythropoietin (EPO), Stem Cell Factor (SCF), and the synthetic glucocorticoid dexamethasone (Dex) (Panzenbock, 1998; von Lindern. 1999). We previously discovered that these culture conditions enabled the essentially unlimited (>10^60^-fold) self-renewal of an immature murine erythroblast derived from transient hematopoietic progenitors in the early mouse embryo (England, 2011). This acquisition of an extensive self-renewal state was accompanied by increased expression of Bmi1 and Ring1a, members of the Polycomb Repressive Complex 1 (PRC1). Importantly, we determined that increasing Bmi1 expression in adult murine erythroid cells with limited self-renewal capacity was sufficient to induce the essentially unlimited self-renewal capacity of their embryonic counterparts (Kim, 2015). These Bmi1-induced self-renewing erythroblasts (iSREs) maintained their capacity to mature not only in vitro but also in vivo where they generated a wave of circulating red blood cells (RBCs) with a normal lifespan (Kim, 2015). BMI1 expression can also expand the self-renewal capacity of adult human-derived erythroblasts (Liu, 2021).

Bmi1 was originally identified as a target of Moloney virus insertion leading to accelerated B lymphoid tumors in mice (van Lohuizen,, 1991). It was subsequently recognized that Bmi1 regulates the self-renewal of normal and malignant stem cells, as well as more mature cell types including fetal-derived B1a lymphocytes (Park, 2003; Kobayashi, 2020). Bmi1 is a member of the polycomb repressive complex 1 (PRC1), which classically functions as a transcriptional repressor through the mono ubiquitination of histone H2A at Lysine 119 (H2AK119Ub) via the E3 ubiquitin ligase, RING1(A/B) (Wang, 2004; Cao, 2005). In this way, Bmi1 can impact multiple cellular functions/pathways including the cell cycle, survival, senescence, and DNA repair (Bhattacharya, 2015). Interestingly, ribosome biogenesis was recognized as a target of Bmi1 in maturing murine erythroblasts, though here Bmi1 directly bound multiple ribosomal protein genes, which was associated surprisingly with gene activation (Gao, 2015). Additionally, recent CUT&RUN studies in erythroid-lineage cells revealed IGF2BP1, IGF2BP3, and Lin28B to be direct targets of BMI1 repression, which in turn regulate fetal hemoglobin levels (Qin, 2023). However, the broader potential functional targets of BMI1 in the erythroid lineage otherwise remain poorly understood.

Here we investigated the function of BMI1 in the regulation of human erythroid self-renewal. Using CUT&RUN as a global and unbiased approach, multiple potential targets of BMI1 were identified including the INK/ARF locus, where BMI1 acts as a repressor to silence the expression of cell cycle inhibitors. Importantly, we also identified a group of upregulated BMI1 target genes not marked by repressive histone modifications following BMI1 overexpression, including key genes regulating cholesterol homeostasis such as HMGCR, the rate-limiting enzyme in cholesterol synthesis. Despite prolonged culture and BMI1 overexpression, iSREs retain the ability to terminally mature with ∼50% rate of enucleation and the ability to agglutinate with typing reagent monoclonal antibodies against conventional Rh, MNS, Kell, and Duffy system antigens. Our findings support the concept that BMI1 acts not only as a transcriptional repressor but also as a transcriptional activator to enhance the in vitro self-renewal of erythroid precursors. Expanding the pool of immature erythroid precursors will ultimately help meet the increasing need for standardized reagent RBCs and for cRBCs for transfusion of alloimmunized patients.

## Methods

### Human biological samples

Peripheral blood mononuclear cells (PBMCs) were isolated from 40 mls of peripheral blood collected from individuals under the University of Rochester School of Medicine IRB approved protocol RSRB00003723. The University of Rochester Institutional Biosafety Committee approved all experiments involved with human peripheral blood. PBMCs, isolated by Ficoll-Paque Premium (Cytiva) according to manufacturer’s instructions, were seeded at 0.5-1 x 10^7^ cells/ml in “phase 1 expansion medium” consisting of Stemspan SFEM (Stem Cell Technologies) supplemented with Stem Cell Factor (SCF; 100 ng/ml, Peprotech), Erythropoietin (EPO; 2 U/ml), water-soluble dexamethasone (1uM, Sigma), IGF-1 (40ng/ml, Peprotech), IL-3 (5 ng/ml, Peprotech), and cholesterol-rich lipids (40 mg/mL, Sigma), and cultured from expansion day 0 to day 5 (Fig. 1A).

**Figure 1.**
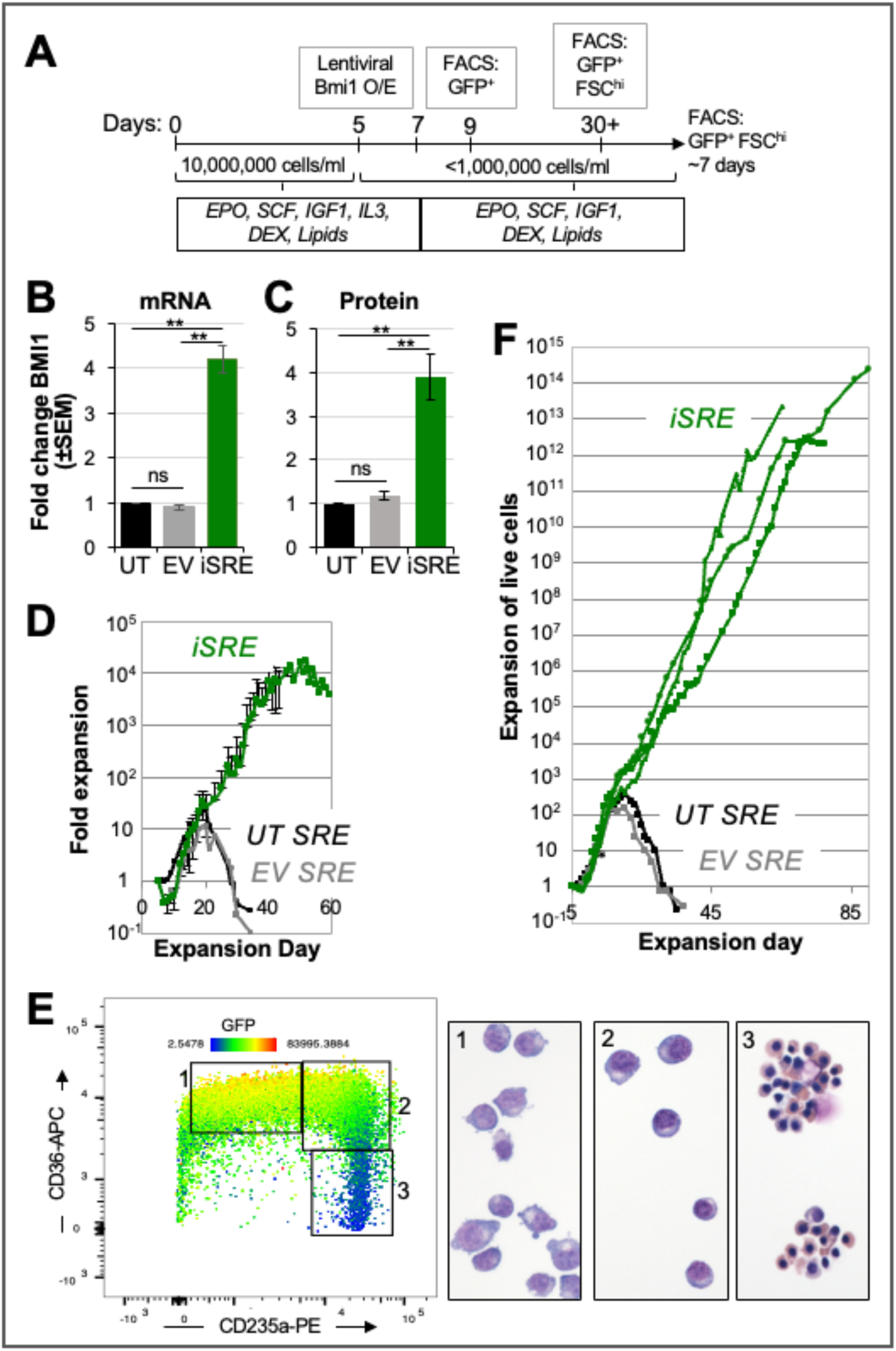
BMI1 expands the self-renewal of human self-renewing erythroblasts (SREs). **(A)** Schema of iSRE establishment and culture from PBMCs. **(B)** Lentiviral transduction leads to an approximately four-fold increase in BMI1 transcript levels compared to compared to untransduced (UT) SREs and empty vector transduced (EV) SREs, which have, as expected, similar levels of BMI1. N=6. p-value was calculated based on paired two-tailed Student t-test. **p<0.05 **(C)** Lentiviral transduction leads to an approximately four-fold increase in BMI1 protein levels compared to compared to UT and EV SREs. N=4. p-value was calculated based on paired two-tailed Student t-test **p<0.01. **(D)** iSREs have approximately a 10^4^-fold expansion of BMI1-induced SREs (iSREs), compared to untransduced (UT) SREs and empty vector transduced (EV) SREs, which have similar limited (200-300-fold) expansion. N=4 independent iSRE cultures. **(E)** FACS-based analysis of iSRE cultures reveals 3 subpopulations of cells. Population 1 is CD36+CD235lo and consist of cells with an immature erythroblast morphology. Population 2 is CD36+CD235+ and consist of cells with a slightly more mature morphology. Population 3 is CD36loCD235+ and consist of smaller hemoglobinizing cells with condensed nuclei. Cells are colored based on GFP expression which is highest in Population 1 and lowest in Population 3. **(F)** Enrichment of immature erythroblasts by weekly fluorescence activated cell sorting (FACS) leads to a 10^12^-fold expansion of BMI1-induced SREs (iSREs), compared to untransduced (UT) and empty vector transduced (EV) control SREs. N=3 independent iSRE cultures.

Human CD34+ cells, purchased from the Yale Cooperative Center of Excellence in Hematology, served as an alternate source of hematopoietic stem and progenitor cells. Cells were cultured for 4 days in IMDM with Penicillin / Streptomycin, 2% PB plasma, 3% AB serum, insulin (10 ug/ml), heparin (3 U/ml), transferrin (200 ug/ml), SCF (10 ng/ml), EPO (3 U/ml), IL-3 (1 ng/ml), and Glutamax (2 mM) (Hu, 2013). The cells were then transferred for 2 days of culture in “phase 1 expansion medium” (Supplemental Fig. 1A).

Human bone marrow cells were isolated from hip surgery samples collected through the Tissue Donation Program at Northwell Health and hip aspirates from the Hospital for Special Surgery, as IRB-approved at Northwell Health, the New York Blood Center and the Hospital for Special Surgery. Primary human proerythroblasts and orthochromatic erythroblasts cells were identified by flow cytometry, according to the immunophenotype described (Yan, 2021).

### Lentiviral constructs and lentiviral transduction

The pReceiver-Lv165 human BMI1 overexpression plasmid (EX-B0015-Lv165, GeneCopoeia) and the pReceiver-Lv165 empty vector plasmid (EX-NEG-Lv165, GeneCopoeia) contain an EEF1A promoter with an IRES2 followed by an eGFP fluorescent marker. Site directed mutagenesis was used to create an additional Lv165 BMI1 overexpression vectors with a P2A sequence instead of the IRES by designing primers using the New England Biolabs (NEB)-base changer website (https://nebasechanger.neb.com) in combination with the Q5 Site-Directed Mutagenesis kit (NEB, E0554S), following the manufacturer’s instructions. Lentiviral particles were harvested from HEK293T 72 hours post transduction, stored at -80C, and added to cultured PBMCs or CD34+ cells at day 5 or day 6, respectively (Fig. 1A and Supplemental Fig. 1A). Following overnight culture, cells were pelleted and resuspended in fresh “phase 1 expansion media” at a concentration of 5x10^5^ cells/ml for 24 hours.

### SRE/iSRE expansion culture

Two or three days following lentiviral transduction, PBMC- or CD34+-derived cells, respectively, were reseeded at 5 x 10^5^ cells/ml in “phase 2 expansion media”, which consists of the same components as “phase 1 expansion media”, but without IL-3 (Fig. 1A and Supplemental Fig. 1A). Following a further two days of culture, DAPI-GFP+ cells were enriched by fluorescence-activated cell sorting (FACS) and resuspended in “phase 2 expansion media” at a concentration of 5X10^5^ cells/ml (Fig. 1A). The DAPI^-^, FSC^hi^, GFP^hi^ self-renewing iSREs were enriched weekly by FACS (Supplemental Fig. 1B). Total live cells were counted daily and the cell concentration was brought to 5x10^5^ total cells/mL with at least half volume medium changes. The daily fold expansion was calculated by the number of live cells counted on that day normalized to the number of live cells that seeded the culture.

To test the function of exogenous factors, iSREs were cultured in “phase 2 expansion media” with the exclusion of either EPO, SCF, or dexamethasone. The minus-dexamethasone condition also included the addition of the glucocorticoid receptor antagonist, RU486 (1μM; Sigma). Total live cells were counted daily.

### iSRE maturation culture

iSRE were transferred at a concentration of 1X10^6^ cells/ml into “maturation media” consisting of 1X Iscove’s Modified Dulbecco’s Medium (IMDM; Gibco), SCF (100 ng/ml), EPO (2 U/mL), 5% plasma-derived serum (PDS; Animal Technologies), 10% serum replacement (Invitrogen), 10% Protein-Free Hybridoma Media (PFHM-II; Invitrogen), 1:10 MTG (12.7 ul/100 ml; Sigma), and 20% BIT (Stem Cell Technologies). The SCF was removed after day 3 of maturation and cells were cultured at 2X10^6^ cells/ml from maturation day 4 to 7.

### RBC gel cards

Cryopreserved iSREs were thawed into “phase 1 expansion media”, cultured for 3 days and resuspended in “maturation media” for 7 days (An, 2022). Matured iSREs were serologically phenotyped for ABO; Rh (D, C, c, E, e); Kell (K, k); Duffy (Fy^a^, Fy^b^); and Glycophorin B (S, s) using a gel-based method (Ortho Diagnostics). Rh antigen typing was performed using commercial monoclonal IgM typing reagents anti-D (Alba Alpha), anti-C, anti-c, anti-E, and anti-e (Immucor Gammaclone) on buffered gel cards (Ortho Clinical Diagnostics). Kell, Duffy, and Glycophorin B antigen typing was performed using commercial anti-K, anti-k, anti-Fya, anti-Fyb, anti-S, and anti-s, (Ortho Sera) on anti-IgG gel cards (Ortho Clinical Diagnostics). Matured iSREs treated with 0.1% ficin solution (Immucor) for 10 minutes at 37 °C were washed with PBS before use. 1 to 1.5 million mature iSREs resuspended in 50 μL of Micro Typing Systems 2 diluent (MTS2, Ortho Clinical Diagnostics) were added to each gel column then incubated with 25 μL of each typing reagent antibody for 30 minutes at room temperature for buffered gel cards or at 37 °C for anti-IgG gel cards before centrifugation on the MTS Workstation (Ortho Clinical Diagnostics) at 1032 rpm for 10 minutes.

Plasma from type O and type A patients with antibody testing performed for clinical care were collected after informed consent under a protocol approved by the IRB at the Children’s Hospital of Philadelphia. 1 to 1.5 million iSREs were added to anti-IgG cards (Ortho Clinical Diagnostics) and incubated with 25 μL plasma at 37 °C for 30 minutes followed by centrifugation on the MTS Workstation (Ortho Clinical Diagnostics).

### Analysis of iSREs

*Morphologic analysis:* 50,000 iSREs in PB2 (Dulbecco phosphate-buffered saline, 0.1% glucose, and 0.3% bovine serum albumin [Gemini Bioproducts]) were cytospun (Shandon II; Thermo-Scientific) onto a glass slide and air dried. Slides were stained with Wright-Giemsa and analyzed using the Nikon Eclipse 80i microscope, DS-Fi1 camera, and NIS Elements Software (Nikon).

*Cell cycle analysis:* BrdU incorporation was performed using the BrdU Flow kit (Supplemental Table 1) according to manufacturer’s instructions. iSREs were stained with BrdU for one hour. Ki67 staining was performed on ethanol-fixed and permeabilized iSRE stained with Ki67-APC antibody (Supplemental Table 1). BrdU and Ki67 data were collected using an LSR-II (BD Biosciences) flow cytometer and analyzed using FlowJo software (version 10; BD Biosciences).

*Apoptosis assay:* 500,000 iSREs were resuspended in Annexin V binding buffer (Fisher, #NC1267456) and stained with Annexin V-AF647 and propidium iodide (Supplemental Table 1) according to manufacturer’s instructions. Data were collected on an LSR-II and analyzed using FlowJo software.

*Karyotype:* iSREs (5x10^6^ cells) from three donors in expansion media were submitted to the URMC Cytogenetics laboratory for G-banding analysis. 20 cells that were in metaphase were counted at a Band Level of 425.

### Flow cytometry

Surface immunophenotyping was carried out on cells blocked in 10% normal mouse serum (Invitrogen) in PB2 and then stained with the antibodies (Supplemental Table 1). DRAQ5 (Ebioscience) was used as a DNA stain when indicated. For lipid analysis, cells were stained with anti-cholera toxin beta unit antibody (CTB, Sigma) according to manufacturer’s instructions, washed in PB2 and stained with surface antibodies as above. For filipin staining, surface stained cells were fixed in 4% formaldehyde for 15 min, washed in PB2 and stained with 100 ug/ml Filipin (Sigma) for 3 hours. Standard flow cytometric analysis was carried out utilizing the LSR-II flow cytometer and FlowJo software. Imaging flow cytometry was carried out using the ImageStream GenX (Amnis/Cytex) and analyzed with the IDEAS software (v6.2, Amnis/Cytex). Cytoplasmic area was determined using a combined Morphology mask based on staining of CD44, CD235a, CD71, CD36, and Adaptive Erode at 81% of the brightfield. These were used alone or in combination with each other depending on the stain used. The nuclear area was determined using the Morphology mask of DNA stain (DRAQ5) mask as previously described (McGrath, 2017).

### Histochemistry

Nile Red staining (Sigma) was performed in solution on paraformaldehyde fixed iSREs, at a concentration of 2.5ug/ml for 30 min at room temperature. Filipin staining (Sigma) also occurred in paraformaldehyde fixed cells at a concentration of 50ug/ml for 2 hr at room temperature after cells were pre-blocked in 1.5mg/ml glycine in PBS. All staining was performed in the dark, and cells were then cytospun on slides, counterstained with DAPI (2ug/ml, Sigma), mounted, and viewed with widefield fluorescence microscopy.

### qPCR

RNA was harvested using the Qiagen RNeasy kit following manufacturer’s guidelines. Quantitative reverse-transcription polymerase chain reaction was performed as described (Kingsley, 2006) using Taqman Gene Expression Assays (Thermofisher, Supplemental Table 2) with an annealing temperature of 60°C. Transcript expression was calculated relative to the 18S control gene.

### Western blotting

Cells were resuspended in RIPA buffer (Cell Signaling Technology), vortexed, sonicated, and frozen. Following 4-20% gradient SDS page gel (Biorad) electrophoresis, proteins were transferred to a 0.2um nitrocellulose membrane and stained with primary antibodies including BMI1 (6964S, Cell Signaling Technology), H4 (13919, Cell Signaling Technology), Beta globin (sc-21757, Santa Cruz), Gamma globin (sc-21756, Santa Cruz), Alpha globin (sc-514378. Santa Cruz), or Epsilon globin (ab156041, Abcam). The HRP-conjugated secondary antibodies were detected with WesternSure PREMIUM Chemiluminescent Substrate per the manufacturer’s instructions and quantitated on a LICOR C-DiGit Blot Scanner.

### RNA-Seq

Sequencing of poly A selected mRNA from three replicates of SRE and iSRE cells was performed at the Yale Center for Genome Analysis. For comparison of iSRE data to stage selected RNA-sequencing data in GEO GSE53983 and GSE128268, reads were mapped to the human genome using STAR https://pubmed.ncbi.nlm.nih.gov/23104886/, and reads in genes were counted using featureCounts https://pubmed.ncbi.nlm.nih.gov/24227677/. For comparison of iSRE and SRE data, reads in genes were quantified using kallisto https://pubmed.ncbi.nlm.nih.gov/27043002/. Differential expression between different sample types was identified using DESeq2 https://pubmed.ncbi.nlm.nih.gov/25516281/.

### CUT&RUN

CUT&RUN was performed as outlined in (Skene, 2017). Proliferating SREs (untransduced) or iSREs in “phase 2 expansion media” were cultured past day 30 and collected for analysis. A minimum of four replicates (two biological and two technical) were performed for each antibody. CUT&RUN was performed using the CUTANA ChIC/CUT&RUN Kit (Epicypher) using 260,000 to 500,000 cells/sample and the following antibodies: BMI1 (D20B7, Cell Signaling), RING1B (D22F2, Cell Signaling), H2AK119Ub (D27C4, Cell Signaling), and H3K27me3 (9733, Cell Signaling).

Libraries were generated using the NEBNext Ultra II DNA Library Prep kit (NEB). For transcription factors (BMI1 and RING1B), size selection was optimized for small fragments (end prep for 20C for 30 minutes followed by 50C for 60 minutes, ligation AMPure XP cleanup at 1.75X, annealing temperature of 65C for library amplification, and library AMPure XP cleanup using 1st round negative selection at 0.8X followed by a 2nd round positive selection at 1.2X). Libraries were sequenced with illumina HiSeq, paired end 150 bp length reads.

Fastq files were aligned to hg38 using Bowtie2, and duplicates were removed using Picard. Read count normalization (RPKM) was performed on alignment files. Replicate variability was assessed by Spearman correlation plots (see supplement) and genome track visualization, resulting in exclusion of one outlier BMI1 iSRE replicate for downstream analyses. Peaks were called on individual replicates using macs2 callpeak (narrowPeak), and union peak sets were used for DiffBind differential analysis in R (BMI1 CUT&RUN iSRE n=5, SRE n=4). Volcano plots were generated in R. Replicate merged bigwig files were generated using bigWigMerge (deepTools) for heatmap and genome track visualization. Heatmaps were generated using plotHeatmap (deepTools). Kmeans clustering was performed with plotHeatmap --kmeans and --outFileSortedRegions. Genomic regions were annotated using Homer annotatePeaks.pl. Region gene associations were determined as single nearest gene within 2kb of TSSs. Enrichr was used to identify gene associations (Chen, 2013).

### Data availability

RNA sequencing data and primary CUT&RUN data are available in GEO at accession GSE253291 and GSE253289, respectively, under superseries GSE253292.

### Statistics

Differences between groups were compared with unpaired one-tailed or two-tailed *Student’s t-test.* A two-tailed paired *Student’s t-test* and two-way ANOVA analyses with Bonferroni correction were used when appropriate and p-values indicated in the figure legends. Statistical analyses were performed using Excel (Microsoft Corp., Redmond, WA) or Prism software (GraphPad Software, San Diego, CA).

## Results

### Increased BMI1 expression drives extensive in vitro expansion of immature human erythroblasts

Adult human erythroblasts normally have a restricted proliferative capacity (Panzenbock, 1998; von Lindern, 1999). Lentiviral transduction of BMI1 on day 5 of PBMCs (Fig. 1A) resulted in an approximately 4-fold increase in both BMI1 transcript and protein levels (Fig. 1B,C and Supplemental Fig. 1C). While untransduced (UT) and empty vector (EV)-transduced control self-renewing erythroblasts (SREs) underwent limited (∼200-400-fold) expansion, BMI1-transduced self-renewing erythroblasts (iSREs) underwent approximately 10^4^-fold expansion (Fig. 1D). Analysis of these cultures at later timepoints of expansion revealed a significant subpopulation of small, maturing cells with condensed nuclei that had downregulated CD36 surface expression, as well as BMI1 expression as evidenced by decreased GFP levels (Fig. 1E, population 3). Since late-stage erythroid precursors can negatively impact the self-renewal of immature erythroblasts (Migliaccio, 2011), we instituted weekly purification of FSC^hi^ GFP^hi^ immature erythroblasts (Supplemental Fig. 1B) by FACS. This approach resulted in approximately 10^12^-fold total expansion of immature human erythroblasts over a 2-3 month period (Fig. 1F). We found similar overall expansion of iSREs when initiated from CD34+ cells (Supplemental Fig. 1D).

### iSRE are poised at an immature erythroblast stage of maturation

iSREs have an immature erythroblast morphology, characterized by a large nuclear:cytoplasmic ratio and nuclear staining suggesting open chromatin (Fig. 2A). Consistent with an immature erythroid phenotype, iSREs are not only CD36+, CD235^mid^ but also CD71+, CD44+, CD49d^lo^, and CD34-(Fig. 2B). The average cell area and nuclear area of self-renewing iSREs remain consistent throughout extensive culture (Fig. 2C). Importantly, despite prolonged culture and increased BMI1 expression, iSREs maintain a normal karyotype and lack evidence of clonal aberrations based on G-banding analysis (Fig. 2D and Supplemental Fig. 1E).

**Figure 2.**
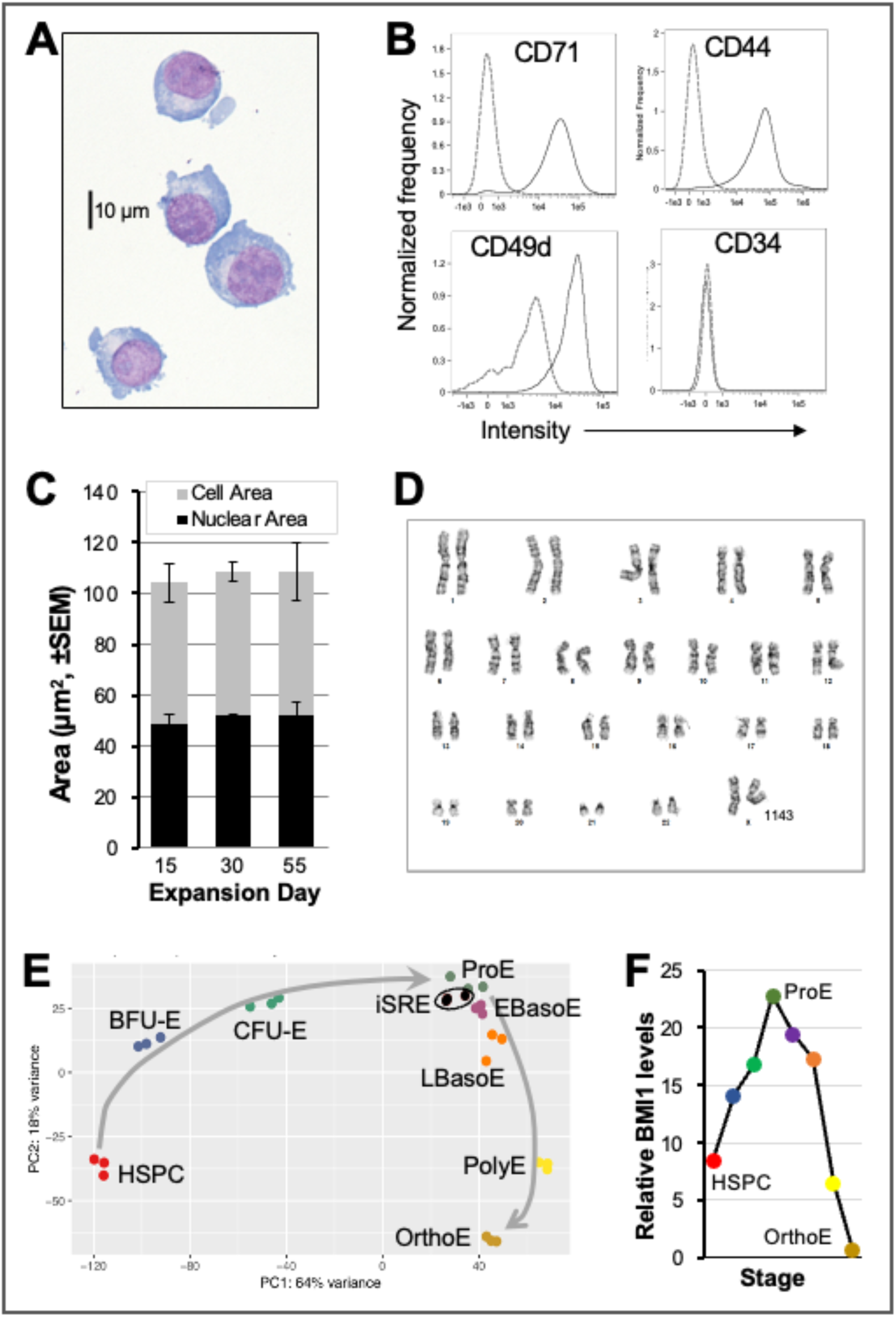
Characterization of human iSREs. **(A)** Morphology of self-renewing iSREs **(B)** Surface expression of CD71, CD44, CD49d, and CD34 on iSREs. **(C)** Consistency of cell and nuclear area of iSREs on days 15, 30, and 55 of culture, as analyzed by imaging flow cytometry. **(D)** Karyotype of iSREs at day 45 of culture. Data from 1 of 3 independent cultures shown. **(E)** Principal Component Analysis (PCA) of global transcriptomes of CD34-derived erythroid populations and of iSREs indicate that iSREs are closest in gene expression identity to proerythroblasts (ProE). HSPC = hematopoietic stem and progenitor cells, BFU-E = burst forming units erythroid, CFU-E = colony forming units erythroid, EBasoE = early basophilic erythroblasts, LBasoE = late basophilic erythroblasts, PolyE = polychromatophilic erythroblasts, and OrthoE = orthochromatic erythroblasts. **(F)** BMI1 transcript levels during the differentiation trajectory of HSPCs to OrthoE.

We compared the transcriptomes of iSREs with that of progressive stages of the human erythron differentiated from CD34+ cells (An, 2014). As shown in Fig. 2E, principal component analysis (PCA) revealed that 3 independent (population 1) iSRE samples cluster together and are found closest to proerythroblasts (ProE). These data, together with the cellular morphology and cell surface phenotype, suggest that SRE/iSRE are poised at a ProE-like stage between upstream erythroid progenitors and downstream terminally maturing erythroid precursors. BMI1 transcript levels increase during erythroid progenitor differentiation, peaking in ProE (Fig. 2F). They subsequently decline very rapidly during terminal maturation of erythroid precursors. This is consistent with the loss of GFP as iSREs spontaneously mature in expansion media (Fig. 1E, population 3).

### iSREs retain the ability to terminally mature in vitro and can serve as reagent red cells

To test if iSREs remain capable of terminal maturation despite prolonged expansion, cells were transferred into maturation media and cultured for 7 days. Morphological analysis revealed a reduction in cell and nuclear size, with acidification of the cytoplasm, and approximately 50% rates of enucleation (Fig. 3A,B; Supplemental Fig. 2A). *BMI1* transcript expression decreases in murine erythroblasts as they terminally mature (Gao, 2015). Consistent with these data and the marked downregulation of BMI1 transcript levels seen in CD34+-derived late erythroblasts (Fig. 2F), we found that iSREs rapidly downregulate BMI1 transcript and protein levels (Fig. 3C; Supplemental Fig. 2B). Maturing iSREs express adult (HBB, HBA) and fetal (HBG) globin genes (Fig. 3D).

**Figure 3.**
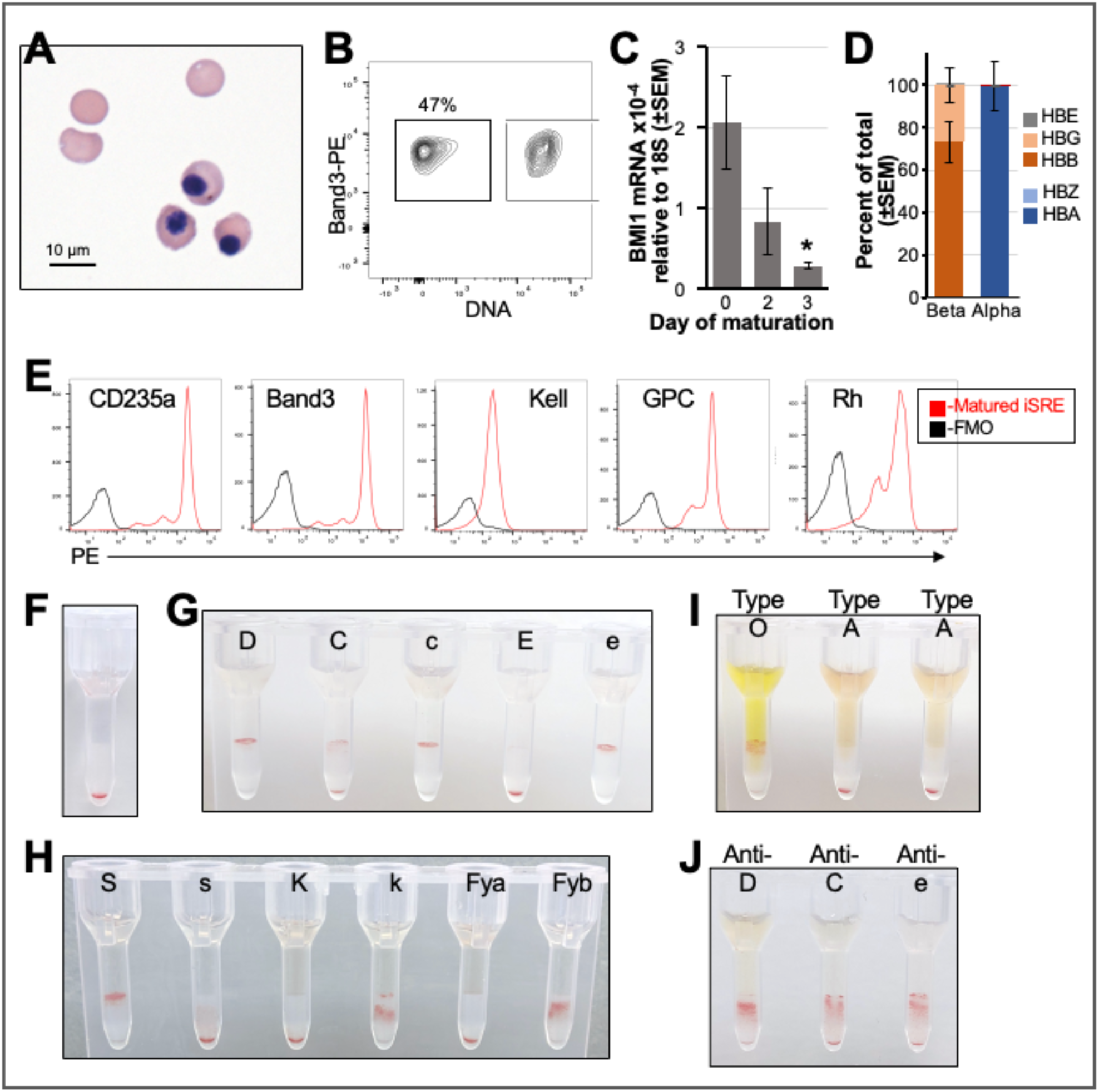
iSREs terminally mature and can serve as reagent red cells. **(A)** Morphology of terminally maturing iSREs at day 8 of maturation culture. **(B)** Flow-based assay of day 8 matured iSREs reveals 47% of erythroid cells have enucleated. **(C)** BMI1 transcript levels rapidly decrease during in vitro maturation of iSREs. N=4. Paired two-tailed Student t-test *p<0.05. **(D)** Globin gene expression of iSREs matured for 3 days and quantified by qPCR. N=3. **(E)** Surface expression of CD235a, Band3, Kell, Glycophorin C (GPC), and Rh on matured iSREs compared to FMO controls. **(F)** Gel card assay of 1.5 million matured iSREs without antigen serving as a negative control. **(G)** Gel card assay of Rh-associated matured iSREs **(H)** Gel card assay of Glycophorin B (S,s), Kell (K,k), and Duffy (Fya,Fyb) systems. **(I)** Gel card assay of iSRE-derived cells agglutinated with plasma from type A and type O donors without any other known antibodies. **(J)** Ficin-treated iSRE-derived cells agglutinated with plasma containing anti-D, anti-C, or anti-e, respectively, from type A patients

We next asked if iSRE-derived cells could be used as an alternative source of reagent RBCs with blood bank assays used to guide donor selection and RBC compatibility (An, 2022). Terminally maturing iSREs are characterized by surface expression of glycophorin A (CD235a), Band3, glycophorin C (GPC), Rh, and Kell (Fig. 3E). To test if iSRE-derived cRBCs had properties of mature erythrocytes (e.g. small size) that are compatible with the standard gel card assay we first added 1.5 million cRBCs to gel columns without addition of reagent antibodies to ensure that the cells could pass through the gel matrix upon centrifugation to form a pellet (Fig. 3F). Agglutination prevents cells from traveling through the gel matrix to the bottom of each column and indicates cell surface expression of the corresponding antigen. iSRE-derived cells agglutinated with typing reagent monoclonal antibodies against conventional Rh antigens D, C, c, and e, demonstrating sufficient Rh expression on the cell surface for agglutination assays (Fig. 3G). These cells also appropriately agglutinated with typing reagent antibodies against the Glycophorin B (S,s), Kell (K,k), and Duffy (Fya,Fyb) systems (Fig. 3H). iSRE-derived cells agglutinated with plasma from a type O donor with expected naturally-occurring anti-A but without any other non-ABO antibodies, and did not agglutinate with plasma from type A donors who lack anti-A (Fig. 3I). Importantly, ficin-treated iSRE-derived cells agglutinated with plasma from group A patients containing anti-D, -C, or -e, respectively (Fig. 3J). Taken together, these data support the concept that iSRE-derived cRBCs could potentially serve as an in vitro source of reagent RBCs for pre-transfusion antibody testing.

### iSRE proliferation remains dependent on BMI1 and exogenous cytokines

Bmi1 expression is necessary for the continued self-renewal of murine self-renewing erythroblasts (England, 2011). We asked if human iSRE self-renewal remains dependent on BMI1 expression. sh-RNA-mediated knock-down of BMI1 transcript levels markedly reduced iSRE proliferation compared to control (Fig. 4A/B). As an alternative approach, we treated iSREs with PTC-209, a small molecule inhibitor of BMI1, which caused dose-dependent reductions in *BMI1* transcript and protein levels (Fig. 4C; Supplemental Fig. 2C,D). PTC-209 treatment resulted in a rapid loss of iSRE proliferation (Fig. 4D). Taken together, these data indicate that BMI1 is necessary for the ex vivo self-renewal of iSREs. Despite overexpression of BMI1, human iSREs, like their murine counterparts (Kim, 2015), remain dependent on the continued presence of EPO, SCF, and dexamethasone for continued self-renewal (Fig. 4E).

**Figure 4.**
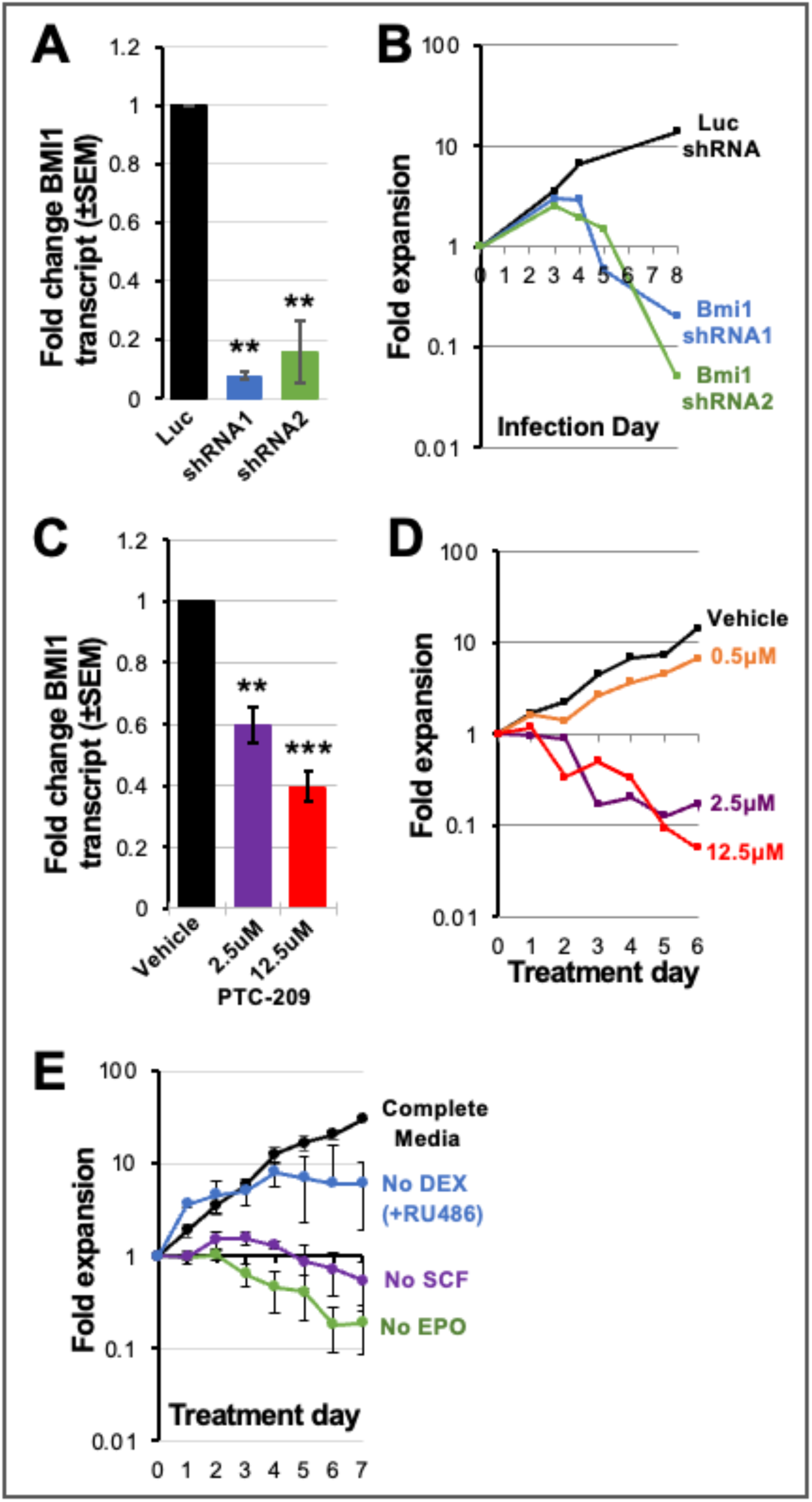
BMI1 regulates the self-renewal of human SREs. **(A)** shRNA knockdown of BMI1 decreases BMI1 transcript levels compared to control luciferase shRNA culture. N=3. p-value was calculated based on one-tailed Student t-test. **p<0.01. **(B)** shRNA knockdown of BMI1 leads to rapid collapse of iSRE cultures. **(C)** PTC-209 inhibition of BMI1 for 24 hours decreases BMI1 transcript levels compared to vehicle-treated controls. N=3. p-value was calculated based on one tailed Student t-test. **p<0.01. ***p<0.001. **(D)** PTC-209 treatment leads to a dose-dependent inhibition of iSRE self-renewal. **(E)** iSRE self-renewal remains dependent on the continued presence of exogenous erythropoietin (EPO), stem cell factor (SCF), and Dexamethasone (Dex).

### Identification of potential downstream targets of BMI1

To gain a better understanding of BMI1 function in erythroid self-renewal we sought to identify direct downstream targets using CUT&RUN to determine genome-wide BMI1 binding sites in both untransduced SREs and BMI1-transduced iSREs. RING1B, a core E3 ubiquitin ligase in PRC1, and its associated histone modification H2A119ub were also included in the analysis because shRNA-mediated knock-down of RING1B significantly impaired expansion of iSREs (Fig. 5A), suggesting a critical role of RING1B in erythroid self-renewal. As dynamic interplay between PRC1 and PRC2 can regulate gene expression, genomic patterns of the PRC2-asociated histone modification H3K27me3 were also determined. A minimum of four replicates were analyzed for each condition and Spearman correlation revealed high agreement between replicates (Supplemental Fig. 3A).

**Figure 5.**
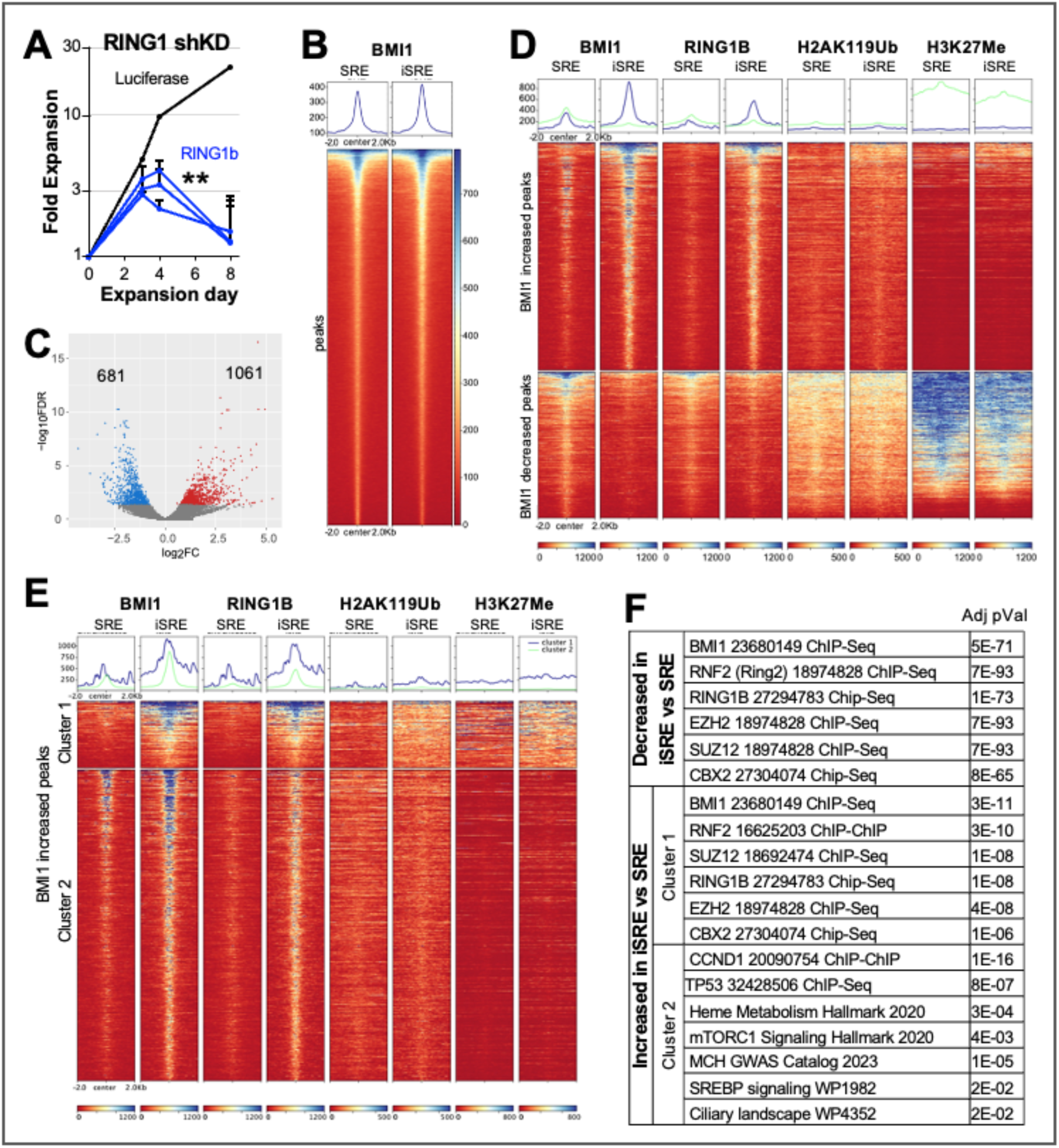
CUT&RUN analyses of BMI1, RING1B, H2A119ub, and H3K27me3 in iSRE and SRE. **(A)** shRNA knockdown of RING1B leads to rapid collapse of iSRE cultures. p-value was calculated based on two-way ANOVA of all RING1B shRNA vs Luciferase. **p<0.01. **(B)** Heatmaps of BMI1 enrichment in untransduced (UT) SREs and iSREs over union peaks (±2 kb). Color scale represents RPKM of each sample using merged replicates (n=4). Sorted based on mean RPKMs. **(C)** Volcano plot showing differential occupancy of BMI1 in iSRE versus untransduced SREs. Significantly increased and decreased peaks are shown in red and blue, respectively, with FDR < 0.05. **(D)** Heatmaps of BMI1 occupancy, RING1B occupancy, H2A119ub, and H3K27me3 over BMI1 differentially increased (top) and decreased (bottom) peaks in untransduced SREs and in iSREs. Color scale represents RPKM of each sample using merged replicates (n=4). Ranking was sorted based on BMI1 in iSREs. Blue line-increased BMI1 peaks. Green line-decreased BMI1 peaks. **(E)** Heatmaps of BMI1, RING1B, H2A119ub, and H3K27me3 over 2 clusters of BMI1 differentially increased peaks (kmeans=2, clustering based on H3K27me3 in iSREs) in untransduced SREs and in iSREs. Color scale represents RPKM of each sample using merged replicates (n=4). Ranking was based on BMI1 in iSREs. Blue line-Cluster 1. Green line-Cluster 2. **(F)** Table of selected gene enrichment terms (Enrichr) significantly (Adjusted p-Value) associated with the indicated groups.

Our CUT&RUN analysis revealed extensive overlap in BMI1 occupancy in SREs and iSREs (Fig. 5B). Consistent with BMI1 overexpression in iSREs, BMI1 enrichment was differentially increased at 1,061 regions compared to SREs (Fig. 5C). Although we found BMI1 differentially decreased at 681 regions, the level of BMI1 changes were more modest compared to regions with increased binding (Fig. 5D, BMI1, green lines). The pattern of RING1B binding overall paralleled BMI1 binding both in SREs and in iSREs (Fig. 5D; Supplemental Fig. 3B). In contrast, global changes in overall H2AK119Ub and H3K27me3 between SRE and iSRE were not striking (Fig. 5D). Of note, regions of decreased BMI1 occupancy had dramatically more enrichment for H2A119ub and H3K27me3 than regions of BMI1 gain (Fig. 5D). The mechanism causing decreased BMI1 binding in iSRE despite increased BMI1 expression is unclear.

However, we speculate that the higher H3K27me3 levels in SRE compared to iSREs in these regions might promote the recruitment of BMI1. Examination of regions with increased BMI1 occupancy in iSREs revealed one subset where H3K27me3 was concordantly increased based on Kmeans clustering (Fig. 5E, Cluster 1). These regions also revealed higher levels of RING1B and H2AK119Ub. Thus, both Cluster 1 and regions with decreased BMI1 occupancy (Fig. 5D, bottom) display the expected association of histone modifications with canonical BMI1 function. Consistent with this, enrichment analysis (Enrichr) of genes associated with these regions revealed a high correlation with genes bound by BMI1 and other components of PRC1, as well as PRC2 components, in published datasets (Fig. 5F; Supplementary Table 3).

In contrast, the larger subset of sites with increased BMI1 occupancy in iSREs was associated with minimal H2AK119Ub and H3K27me3 marks (Fig. 5E, Cluster 2). Gene enrichment analysis of Cluster 2 did not reveal strong correlations with known PRC1/2 data sets but did show enrichment of genes known to be regulated by BMI1 (e.g., CCND1, p53) and, interestingly, processes associated with erythropoiesis (e.g., heme metabolism, MCH GWAS) (Fig. 5F, Supplementary Table 3). RING1B was also increased in Cluster 2 iSREs in contrast to H2AK119ub (Fig. 5E, green line), consistent with RING1B’s function in some contexts independently of its ubiquitin ligase activity (Boyle, 2020; Eskeland, 2010; Francis, 2004; Zhang, 2021). Taken together, Cluster 2 findings suggest that acquired BMI1 is acting through mechanisms other than H2AK119Ub deposition.

Approximately half (53%) of all differentially increased BMI1-occupied regions are located at promoters, while the differentially decreased BMI1-occupied sites are located at promoters (27%), introns (33%), and intergenic regions (19%) (Fig. 6A; Supplemental Fig. 3C). We found that BMI1 was enriched near (i.e., within 2kb of) the transcription start sites (TSS) of 6,538 genes in iSREs (Fig. 6B, Supplemental Table 3). Paralleling our genome-wide BMI1 occupancy findings, increases in BMI1 enrichment at TSSs was strongly associated with increases in RING1B, whereas the associations between BMI1 and H2AK119Ub or H3K27me3 were not as strong (Fig. 6B; Supplemental Fig. 3D). Moreover, despite the increase in occupancy of the ubiquitin ligase RING1B, there was no overall increase in enrichment of H2A119ub at promoters in iSRE, further suggesting that BMI1 overexpression can alter gene expression via mechanisms other than H2A119ub deposition.

**Figure 6.**
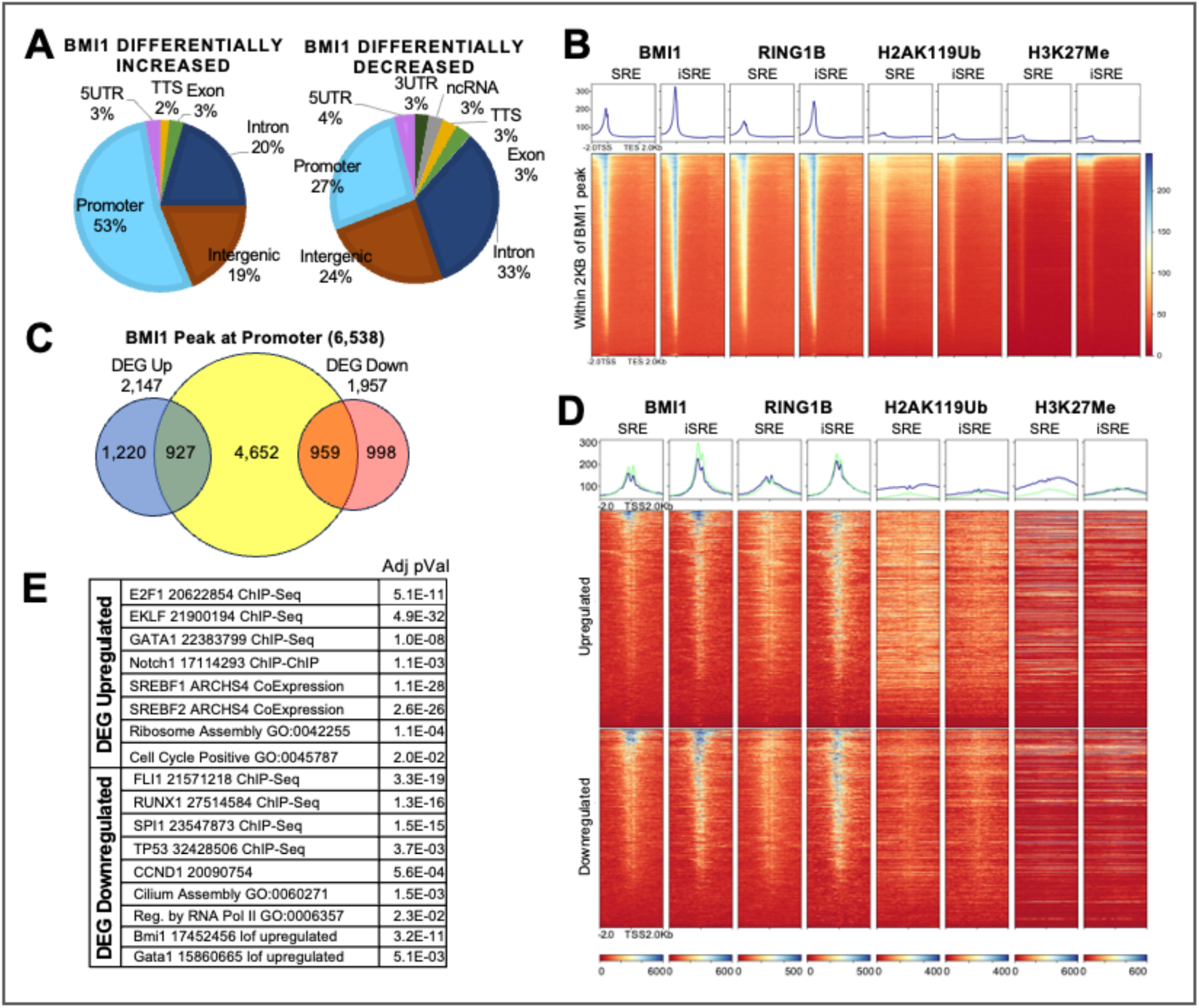
CUT&RUN analyses of BMI1, RING1B, H2A119ub, and H3K27me3 in iSRE and SRE. **(A)** Genomic annotations of BMI1 differentially increased peaks (left) and decreased peaks (right). **(B)** Heatmaps of BMI1, RING1B, H2A119ub, and H3K27me3 associated with promoters in iSRE (genes scaled TSS to TES ± 2kb) in untransduced SREs and iSREs. Color scale represents RPKM of each sample using merged replicates (n=4). Ranking was based on BMI1 in iSREs. **(C)** Venn Diagram showing overlap in BMI1-occupied promoters and differentially upregulated (blue) and downregulated (red) genes. **(D)** Heatmaps of BMI1, RING1B, H2A119ub, and H3K27me3 over promoters of differentially expressed, upregulated (top) and downregulated (bottom), genes (TSS +/-2kb) in untransduced SREs and in iSREs. Color scale represents RPKM of each sample using merged replicates (n=4). Ranking was based on BMI1 in iSREs. Blue line-upregulated genes. Green line-downregulated genes. **(E)** Table of selected gene enrichment terms (Enrichr) significantly (Adjusted p-Value) associated with differentially upregulated and downregulated genes that have BMI1 associated with their promoters.

To further examine potential targets modulated by BMI1 in iSRE, we compared the global transcriptome of iSREs with that of empty vector transduced SREs. There was high concordance of all samples with highest concordance among replicates (Supplemental Fig. 3E). Furthermore, we identified 2,147 up- and 1,957 down-regulated genes (Supplemental Fig. 3F; Supplemental Table 3). Consistent with the well-established role of BMI1 as a transcriptional repressor, nearly half of the differentially downregulated genes were bound by BMI1 (Fig. 6C). Of note, 43% (927 of 2,147) of upregulated genes were also bound by BMI1 (Fig. 6C). Consistent with these findings, BETA analysis, which integrates transcriptomic and protein binding/epigenetic data, strongly predicted that BMI functions to activate, as well as repress, gene expression (Supplemental Fig. 3G) (Wang, 2013). The levels of BMI1 and RING1B occupancy in iSREs were also similarly increased both in upregulated and in downregulated genes (Fig. 6D, blue and green lines, respectively), while the majority of downregulated genes did not accumulate the repressive chromatin H2AK119Ub and H3K27me3 marks (Fig. 6D; Supplemental Fig. 3H).

PRC1 and PRC2 occupancy, as well as H2A119ub deposition, have been associated with CpG islands (Blackledge, 2021; Kallin ‘09). In erythroid lineage cells, BMI1 occupancy at CpG islands was identified at the Lin28b, IGF2BP1 and IGF2BP3 genes, where it was associated with gene repression (Qin, 2023). Consistent with this report, we found that BMI1, RING1B, as well as H2A119ub and H3K27me3 were localized at the CpG islands of these 3 genes, whose expression was repressed, particularly in iSREs (Supplemental Fig. 4A,B). To further investigate the relationship of BMI1 and CpG island occupancy in iSREs, we undertook a global analysis and found that BMI1 and RING1B occupancy were higher at genes whose promoter contained CpG islands than those that did not contain CpG islands (Supplemental Fig. 4C, comparison of CpG at TSS vs No CpG). However, as seen with BMI1 and RING1B occupancy, BMI1 binding at CpG islands was associated equivalently with both upregulated and downregulated genes (Supplemental Fig. 4C, dark blue and green lines, respectively).

Gene enrichment analysis of genes downregulated in iSREs compared to SREs revealed associations with transcription factors of myeloid and megakaryocyte lineages (FLI1, RUNX1, SPI1), as well as regulators of cell cycle and apoptosis (Fig. 6E). Downregulated genes also overlapped with gene sets upregulated by BMI1 loss of function in other cell types (e.g., TP53, CCND1). In contrast, upregulated genes revealed correlations with known regulators of erythropoiesis (e.g., GATA1, EKLF) and cholesterol and lipid homeostasis (e.g., SREBF1, SREBF2), as well as gene sets associated with known BMI1 functions including positive regulation of the cell cycle and ribosomal proteins (Fig. 6E; Supplemental Table 4). Taken together, our CUT&RUN data suggest roles for BMI1 in regulating self-renewal in erythroblasts that extend beyond the classical repressive function ascribed to PRC1, and reveal potentially novel pathways upregulated by BMI in iSRE, including cholesterol metabolism.

### BMI1 occupies the INK/ARF locus and regulates the cell cycle of iSREs

To begin investigating direct targets of BMI1 in erythroid self-renewal, we focused first on the INK/ARF locus, a known repressive target of BMI1 (Jacobs, 1999; Dhawan, 2009). We confirmed BMI1 and RING1B occupancy and H2A119ub deposition specifically at CDKN2A, as well as CDKN2B, which encode the pl6^1NK4a^ and pl4^ARF^, as well as p15^INK4B^ (Fig. 7A). Our CUT&RUN studies also revealed that iSRE had an increase in RING1B occupancy and H2A119ub compared to SRE at these loci. Consistent with the repressive function of PRC1, the increased BMI1 expression in iSREs was associated with reduced expression of p16, p14, and p15 in iSREs compared to SREs (Fig. 7B).

**Figure 7.**
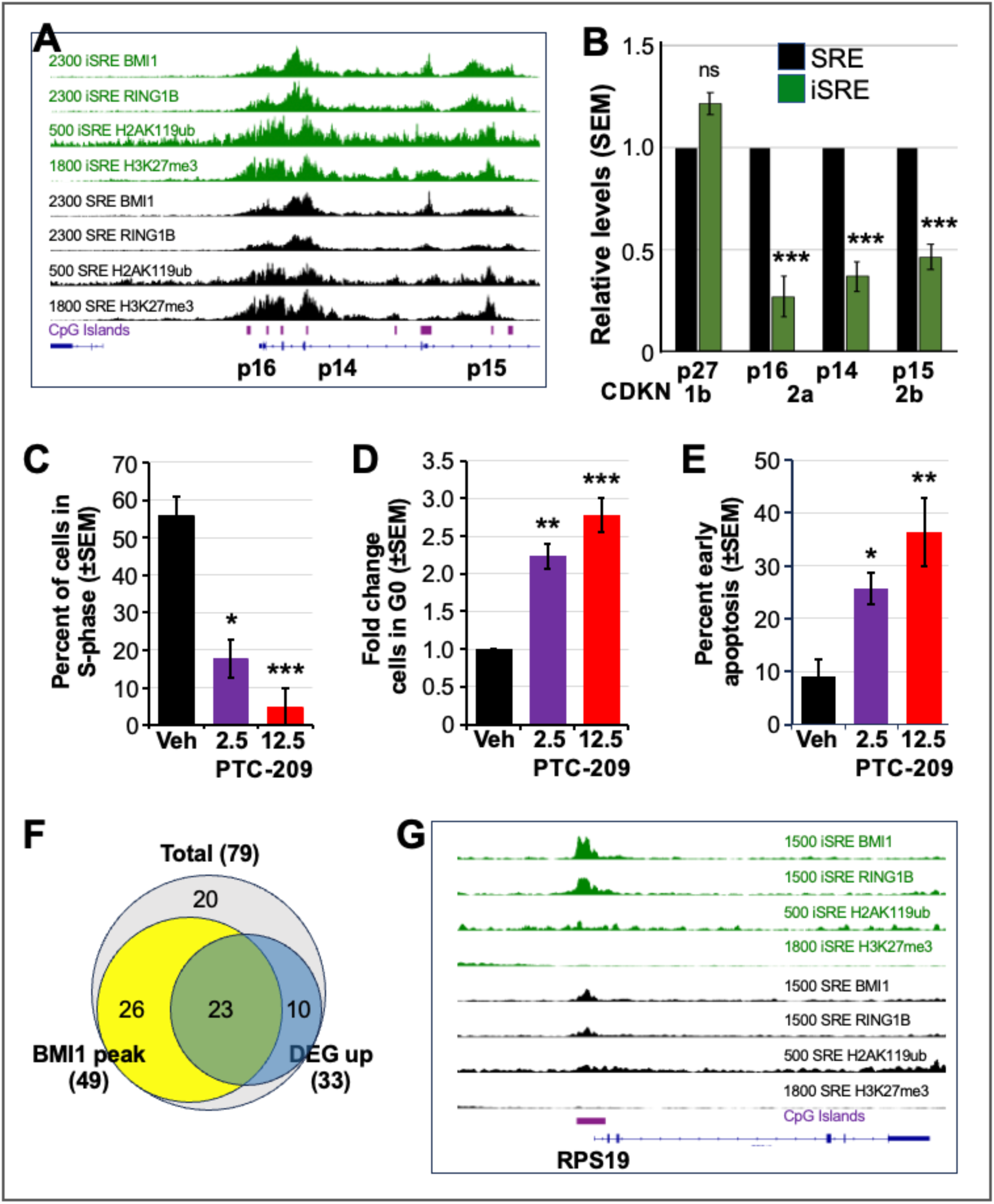
BMI1 regulates the cell cycle in iSREs. **(A)** Occupancy of indicated factors and presence of CpG islands in SRE (black) and iSRE (green) at the INK/ARF locus. Data represent merged replicates (n=4) for all CUT&RUN studies. **(B)** The INK/ARF locus-associated genes are expressed at lower levels in iSREs compared to SREs. N=6. p-value was calculated using a two-tailed Student t-test.***p<0.001. **(C)** The BMI1 inhibitor PTC-209 treatment for 12 hours decreases the percentage of iSREs in S-phase (BrdU analysis) in a dose-dependent manner compared to vehicle (Veh) treated controls. N=3. p-value was calculated using a two-tailed Student t-test. *p<0.05. ***p<0.001. **(D)** PTC-209 treatment of iSREs for 48 hours leads to a dose dependent increase of cells in G0 phase of the cell cycle. N=5. p-value was calculated based on two-tailed Student t-test. **p<0.01. ***p<0.001. **(E)** PTC-209 treatment for 12 hours leads to a dose dependent increase in early apoptotic (Annexin V-positive, PI-negative) cells. N=4. Two-tailed Student t-test *p<0.05. **p<0.01. **(F)** Venn Diagrams showing overlap in BMI1 occupied ribosomal protein genes (yellow) and ribosomal protein genes differentially upregulated in iSREs vs. SREs (blue). **(G)** Occupancy of indicated factors in SRE (black) and iSRE (green) and CpG islands (purple) at the Ribosomal Protein S19 (RPS19) locus. Data represent merged replicates (n=4) for all CUT&RUN studies.

Having identified BMI1 binding at the INK/ARF locus, we more directly tested the function of BMI1 in the regulation of the cell cycle. Treatment of iSREs with the BMI1 inhibitor PTC-209 caused a dose-dependent loss of S-phase (BrdU+) cells at 12 hours post treatment (Fig. 7C). Cell cycle analysis also revealed a significant increase of cells in G0 (Ki67-) at 48 hours of PTC-209 treatment (Fig. 7D). These studies also revealed an increase in sub-G0 cells (Supplemental Fig. 5D), consistent with an increase in cell death of iSREs. p14ARF, which is bound by BMI1, regulates apoptosis, and p14 expression is downregulated in iSREs compared to SREs (Fig. 7B). We therefore analyzed rates of apoptosis in iSREs following BMI1 inhibition. As shown in Fig. 7E, PTC-209 induced a dose-dependent increase in early apoptotic (PI-, Annexin5+) cells. Taken together, these findings support the hypothesis that iSRE self-renewal is controlled in part by BMI1-mediated regulation of cell cycle and cell survival via the INK/ARF locus.

### BMI1 positively regulates expression of ribosomal protein genes in iSREs

Our Gene Enrichment analyses revealed positive regulation of ribosomal proteins (Fig. 6E). Given that ribosomal protein genes have been reported to be bound by Bmi1 in murine erythroid precursors (Gao, 2015), we next examined BMI1 occupancy of these genes in human iSREs. BMI1 and RING1B were bound to the promoters of 49 of 79 ribosomal protein genes (Fig. 7F), as exemplified by the RPS19 gene (Fig. 7G). However, in contrast to the INK/ARF locus, there was minimal H2A119ub or H3K27me3 deposition at these loci. Consistent with the lack of repressive marks, 23 of the 49 ribosomal protein genes bound by BMI1 were upregulated in iSREs compared to SREs (Fig. 7F). These data, taken together with our findings at the INK/ARF locus above, highlight contrasting roles for BMI1 and RING1B-both the classical repressive function ascribed to PRC1, as well as binding of positively regulated genes in the absence of repressive marks.

### BMI1 regulates cholesterol homeostasis in iSREs

Our CUT&RUN studies also revealed that BMI1 binds the gene promoter regions of the rate limiting enzymes of cholesterol biosynthesis (HMGCR, SQLE), of other enzymes in the cholesterol biosynthesis pathway (HMGCS1), of the primary cholesterol importer gene (LDLR), of the genes driving cholesterol esterification (ACAT1, ACAT2), as well as of the primary transcriptional regulator of cholesterol synthesis (SREBP2) (Fig. 8A, asterisks), Fig. 8B, Supplemental Fig. 6). Similar to our findings of RPS19 and other ribosomal protein genes (Fig. 7F,G), CUT&RUN studies also revealed minimal H2A119ub and H3K27me3 marks despite BMI1 and RING1B occupancy at these loci, suggesting a lack of PRC1-mediated repression (Fig. 8B, Supplemental Fig. 6). Consistent with this notion, treatment of iSRE with the BMI1 inhibitor PTC-209 resulted in decreased expression of HMGCR and HMGCS1, while the upregulation of p16 in the same iSREs served as a control for the activity of the BMI1 inhibitor (Fig. 8C).

**Figure 8.**
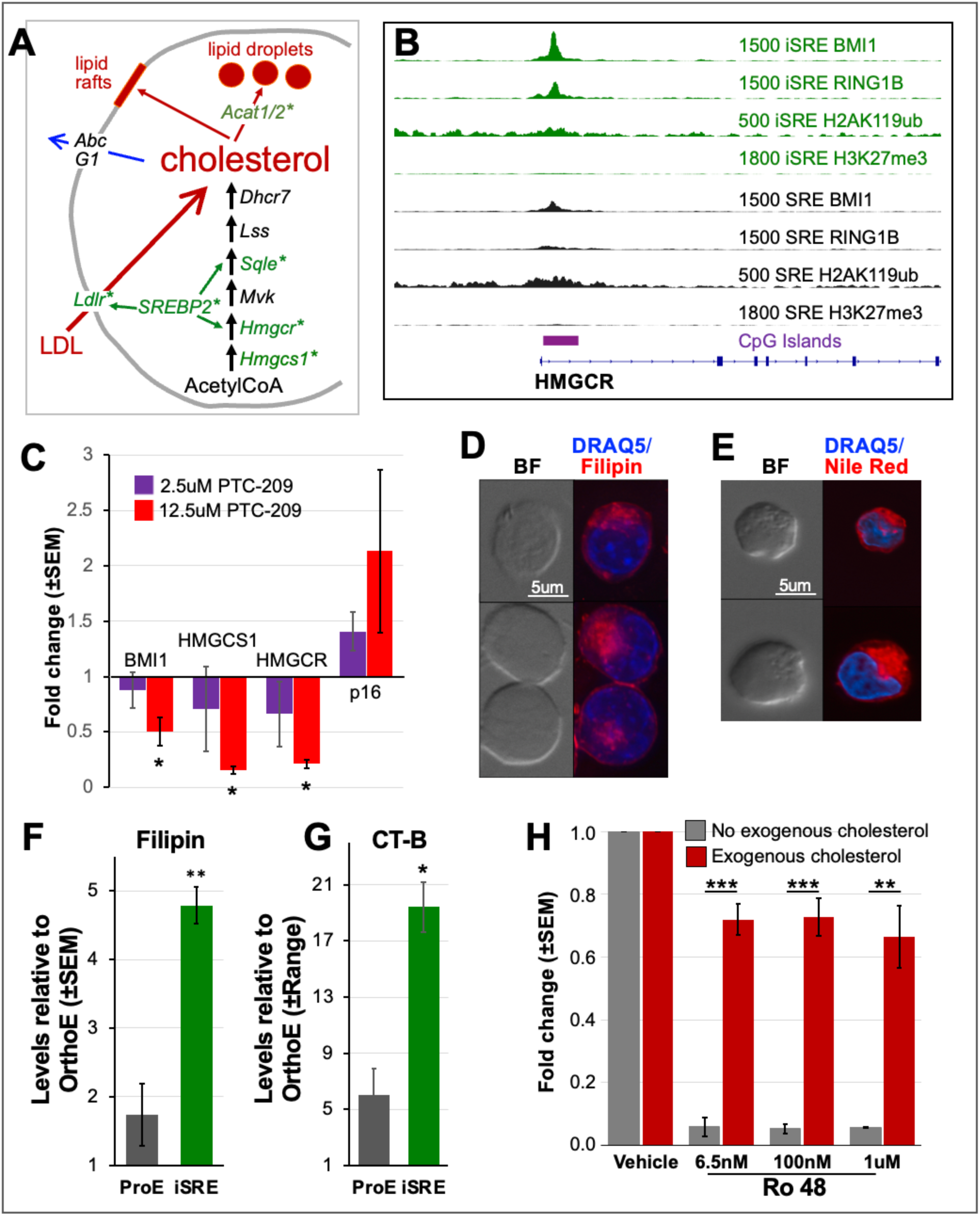
BMI1 regulates cholesterol homeostasis. **(A)** Cholesterol homeostasis depends on cholesterol synthesis, import, export and storage. **(B)** CUT&RUN studies reveal BMI1 and RING1B bind the HMGCR locus at higher levels in iSRE than SRE. Data represent merged replicates (n=4) for all CUT&RUN studies. **(C)** The BMI1 inhibitor PTC-209 decreases the expression of BMI1, HMGCR, and HMGCS1 in iSREs, while increasing expression of p16. N=6. p-value was calculated using a two-tailed Student t-test. *p<0.05 **(D)** iSREs contain cytoplasmic droplets that stain with filipin (cholesterol). **(E)** iSREs contain cytoplasmic droplets that stain with Nile Red (lipids). **(F)** iSREs contain higher levels of cholesterol (filipin stain) compared to primary human proerythroblast (ProE). Staining of both cell populations normalized to primary orthochromatic erythroblasts (OrthoE). N=3 ProE, N=4 iSRE. Two-tailed Student t-test. **p<0.01. **(G)** iSREs contain higher levels of lipid rafts (CT-B staining) compared to primary human ProE. Staining of both cell populations normalized to primary OrthoE. N=2. Error bar indicates range. Paired two-tailed Student t-test. *p<0.05. **(H)** While short-term removal of exogenous lipids does not alter iSRE proliferation, concomitant block of cholesterol synthesis with Ro 48 obviates iSRE proliferation, which can be partially rescued by addition of exogenous lipids. N=3. p-value was calculated based on two-tailed Student t-test. **p<0.01, ***p<0.001.

We next examined the accumulation of cholesterol in proliferating iSREs using an immunohistochemical approach with Filipin and Nile Red staining of cholesterol and lipids, respectively. Both stains revealed accumulation of lipids not only in the cell membrane but also in lipid droplets (Fig. 8D/E), consistent with the vacuoles present in the cytoplasm of iSRE (Fig. 2A). Since primary human erythroblasts lack such vacuoles, we compared the total levels of Filipin staining in iSREs to primary human erythroid cells isolated from the bone marrow. Filipin staining was approximately 1.7-fold higher in ProE than orthochromatic erythroblasts (OrthoE), while iSRE contain approximately 2.8-fold higher levels than marrow ProE (Fig. 8F). Since membrane cholesterol can be localized to lipid rafts, we used Cholera toxin subunit B (CT-B) staining to quantify lipid rafts and found approximately 3-fold higher levels of staining in iSREs compared to normal marrow ProE (Fig. 8G). Talen together, these data indicate that iSREs accumulate cholesterol, which localizes in lipid droplets and in lipid rafts. These data also indicate that cholesterol content and lipid raft levels normally decrease as ProE undergo terminal differentiation, which also parallels the decrease seen in BMI1 transcript and protein levels.

To test the function of cholesterol synthesis and import on iSRE self-renewal, iSREs were cultured for 48 hours in the absence of cholesterol-rich lipids normally added to our cultures. This short-term removal of exogenous lipids did not affect iSRE proliferation (Fig. 8H, vehicle). However, concomitant treatment of the cultures with increasing concentrations of the cholesterol synthesis inhibitor, Ro 48, led to near complete cessation of proliferation (Fig. 8H). Addition of exogenous cholesterol-rich lipids partially rescued iSRE self-renewal. The incomplete rescue suggests that iSREs depend both on cholesterol import and on endogenous cholesterol synthesis to maintain their proliferative capability.

## Discussion

RBCs are the most common form of cell-based therapy with approximately 10 million units transfused annually in the US and over 110 million worldwide. Cultured (c)RBCs could provide reagent red cells for pre-transfusion testing and ultimately supplement donor-derived units (An, 2022), particularly for alloimmunized patients requiring chronic transfusions for whom compatible units are difficult to obtain. However, the limited proliferative capacity of erythroid progenitors and the fixed number of cell divisions associated with terminal erythroid precursor maturation is a major obstacle to generate sufficient numbers of cRBCs for clinical purposes.

Increasing the self-renewal capacity of an erythroid progenitor would facilitate the production of greater numbers of cRBCs from the limited number of starting stem and progenitor cells available in the peripheral blood even after mobilization.

Here, we confirmed that increasing the expression of BMI1 in erythroid progenitors leads to an expansion in the self-renewal capacity of human erythroid cells (Liu, 2021). Four-fold overexpression of BMI1 was sufficient to increase the ex vivo self-renewal capacity of human erythroblasts by 10 billion-fold compared to untransduced and empty vector control cultures.

Morphology, immunophenotyping, and global transcriptomic studies indicate that human iSREs share most similarity with proerythroblasts, which is the stage of erythropoiesis where BMI1 levels normally peak (Fig. 2E). We used a non-biased approach to screen for potential downstream targets of BMI1 in human iSREs to better understand the function of BMI1 in erythroid self-renewal.

Consistent with BMI1 regulation of the cell cycle in multiple cell types (Calés, 2008; Rizo, 2009; Kobayashi, 2020), we found that BMI1 and RING1B each occupy the INK/ARF locus in SREs and iSREs, which was associated with H2A119ub and H3K27Me3 deposition. These data exemplify the canonical repressive activity associated with BMI1-containing polycomb repressive complexes. Functional studies provide data consistent with the notion that BMI1 enhances the cell cycle and inhibits apoptosis in iSREs potentially expanding their self-renewal capacity.

While BMI1, as a member of PRC1 along with RING proteins, is thought to act predominantly in gene repression, our CUT&RUN and transcriptomic studies surprisingly revealed nearly as many upregulated as downregulated genes associated with BMI1 occupancy in iSREs (Fig. 6C). A role for BMI1 in gene upregulation is consistent with studies in murine erythroblasts (Gao, 2015) demonstrating that ribosomal proteins, such as RPS19, were both bound and upregulated by BMI1, as well as a recent study in spermatogonia where BMI1 was also found to act as a transcriptional activator (Liu, 2023). Despite extensive RING1B – BMI1 co-occupancy, the upregulated genes were in general not associated with H2A119ub and H3K27me3 deposition.

RING1B is a relatively weak ubiquitin ligase, particularly when in a complex with BMI1 (Taherbhoy, 2015), and can regulate gene expression independently of its ubiquitin ligase activity (Boyle, 2020; Eskeland, 2010; Francis, 2004; Zhang 2021) Together these data suggest that BMI1 regulates gene expression via other mechanisms than canonical PRC1 activity.

Our CUT&RUN studies also revealed cholesterol homeostasis as a novel and direct target of BMI1. The levels of intracellular cholesterol are dynamically balanced by several processes including 1) synthesis, regulated by the master transcription factor SREBP2, 2) import primarily from LDL via LDLR also regulated by SREBP2, 3) export via ABC transporters, and 4) storage via esterification by ACAT1 and ACAT2 and transport into and out of lipid droplets. Recently, downregulation of cholesterol homeostasis was shown to regulate terminal erythropoiesis (Lu, 2022). However, highly proliferating cells, including many cancer cell types, require high levels of cholesterol (reviewed by Huang, 2020). In addition, high erythroid output disorders, such as hereditary spherocytosis and thalassemia intermedia, as well as polycythemia vera are associated with hypocholesterolemia, suggesting that expansion of erythropoiesis leads to increased utilization of serum cholesterol (Gilbert, 1981; Shalev, 2007). Dexamethasone has been shown to alter lipid metabolism in human erythroblasts making them dependent on import of exogenous lipids via LDL and VLDL (Zingariello, 2019). Consistent with these findings, we found that removal of exogenous lipids from the culture media reduced human erythroblast self-renewal. However, this effect only occurred after several days of culture, suggesting that iSREs can compensate through the use of stored cholesterol and/or by de novo synthesis. Our studies furthermore suggest that iSREs depend both on cholesterol import and on cholesterol synthesis to support their proliferation (Fig. 8H).

We found that BMI1 occupied multiple gene loci associated with cholesterol synthesis, import, and esterification, suggesting extensive regulation of cholesterol homeostasis. Furthermore, upregulated genes bound by BMI1 in iSREs were enriched for genes associated with lipid metabolism and SREBP2. Consistent with these findings, iSREs accumulate large amounts of cholesterol, which is localized not only in the cell membrane associated with increased lipid rafts but also in an extensive network of lipid droplets. BMI1 has previously been implicated in regulating cholesterol homeostasis in glioblastoma cells, however, in these tumor cells BMI1 was considered to be a negative regulator of cholesterol synthesis via an indirect and undefined pathway (Freire-Beneitez, 2021). In contrast, BMI1 appears to act in iSREs as a positive regulator of cholesterol biosynthesis. This includes directly binding to the promoters and increasing the expression of the regulators of the first 2 steps of cholesterol biosynthesis, including the rate-limiting HMGCR.

Despite overexpression of BMI1, which can contribute to immortalization of multiple different cell types (Dimri, 2002; Song, 2006; Tan, 2020), iSREs maintain a normal karyotype and a strong tendency to terminally differentiate, necessitating weekly FACS-based enrichment to maintain a population of self-renewing cells. Our transcriptomic analysis indicates that iSREs appear poised between upstream proliferating erythroid progenitors and downstream terminally maturing erythroid precursors. Indeed, transfer of iSREs into maturation media leads to the rapid downregulation of BMI1 and their terminal differentiation into orthochromatic erythroblasts and reticulocytes, with approximately 50% rates of enucleation. We tested the ability of these differentiated cells to serve as reagent red cells. Using standard gel card-based assays iSRE-derived cells correctly typed Rh, S/s, Kell and Duffy antigens. BMI1-driven expansion of the pool of immature erythroid precursors coupled with future improvements in terminal maturation provide a basis for the eventual generation of sufficient numbers of cultured RBCs to study of erythroid intrinsic disorders and to use clinically for RBC antibody identification, drug delivery, and transfusion therapy.

## Acknowledgements

We acknowledge the diligent work of Jayme L. Olsen, who established the culture of human SREs and iSREs in the lab, characterized the proliferating and terminally maturing iSREs, analyzed the role of the INK/ARF locus and cholesterol in erythroid self-renewal, and wrote early drafts of this paper, but is unfortunately unavailable to approve this version of the manuscript. We acknowledge the assistance of the URMC Flow Cytometry Shared Resource. Funding sources include NIH U01 HL134696 (S.T.C, J.P., C.M.W.), NIH R01 HL130670 (J.P.), NIH R01 DK124777 (L.A.S), NIH R01 DK104920 (L.S.), NIH R01 HL144436 (L.B.), and NIH UR CTSI pilot funding (JLO). We also acknowledge the Yale Cooperative Center of Excellence in Hematology (NIH U54 DK106857) from which we purchased frozen CD34+ cell aliquots.

## Author contributions

K.E.M, K.M., V.P.S., and H.H.A. designed and conducted experiments, analyzed data, and wrote the manuscript.

Ju.P. provided human bone marrow cells.

A.D.K., M.G., A.R.K., B.Z., T.L.S., P.D.K., and C.M.W. designed and conducted experiments.

L.B. and P.G.G. interpreted data and wrote the manuscript.

S.T.C., L.A.S., and J.P. designed experiments, analyzed data, and wrote the manuscript.

## References

An X, Schulz VP, Li J, et al. Global transcriptome analyses of human and murine terminal erythroid differentiation. Blood 2014;123(22):3466–3477.

An HH, Gagne AL, Maguire JA, et al. The use of pluripotent stem cells to generate diagnostic tools for transfusion medicine. Blood 2022;140(15):1723–1734.

Bhattacharya R, Mustafi SB, Street M, Dey A, Dwivedi SK. BMI-1: At the crossroads of physiological and pathological biology. Genes Dis 2015;2(3):225–239.

Boyle S, Flyamer IM, Williamson I, Sengupta D, Bickmore WA, Illingworth RS. A central role for canonical PRC1 in shaping the 3D nuclear landscape. Genes Dev 2020;34(13-14):931–949.

Bryder D, Rossi DJ, Weissmanm IL. Hematopoietic stem cells: the paradigmatic tissue-specific stem cell. Am J Pathol 2006;169(2):338–346.

Calés C, Román-Trufero M, Pavón L, et al. Inactivation of the polycomb group protein Ring1B unveils an antiproliferative role in hematopoietic cell expansion and cooperation with tumorigenesis associated with Ink4a deletion. Mol Cell Biol 2008;28(3):1018–28.

Cao R, Tsukada Y, Zhang Y. Role of Bmi-1 and Ring1A in H2A ubiquitylation and Hox gene silencing. Mol. Cell 2005;20(6):845–854.

Chen EY, Tan CM, Kou Y, et al. Enrichr: interactive and collaborative HTML5 gene list enrichment analysis tool. BMC Bioinformatics 2013;14:128.

Dhawan S, Tschen SI, Bhushan A. Bmi-1 regulates the Ink4a/Arf locus to control pancreatic beta-cell proliferation. Genes Dev 2009;23(8):906–911.

Dimri GP, Martinez JL, Jacobs JJ, et al. The bmi-1 oncogene induces telomerase activity and immortalizes mammary epithelial cells. Cancer Res 2002;62(16):4736–4745.

England SJ, McGrath KE, Frame JM, Palis J. Immature erythroblasts with extensive ex vivo self-renewal capacity emerge from the early mammalian fetus. Blood 2011;117(9):2708–2717.

Eskeland R, Leeb M, Grimes GR, et al. Ring1B compacts chromatin structure and represses gene expression independent of histone ubiquitination. Mol Cell 2010;38(3):452–64.

Francis NJ, Kingston RE, Woodcock CL. Chromatin compaction by a polycomb group protein complex. Science 2004;306(5701):1574-1577.

Freire-Beneitez V, Pomella N, Millner TO, et al. Elucidation of the BMI1 interactome identifies novel regulatory roles in glioblastoma. NAR Cancer 2021;3(1):zcab009.

Gao R, Chen S, Kobayashi M, et al. Bmi1 promotes erythroid development through regulating ribosome biogenesis. Stem Cells 2015;33(3):925–938.

Gilbert HS, Ginsberg H, Fagerstrom R, Brown WV. Characterization of hypocholesterolemia in myeloproliferative disease. Relation to disease manifestations and activity. Am J Med 1981;71(4):595–602.

Hu J, Liu J, Xue F, et al. Isolation and functional characterization of human erythroblasts at distinct stages: implications for understanding of normal and disordered erythropoiesis in vitvo. Blood 2013:121(16):3246–3253.

Huang B, Song B, Xu C. Cholesterol metabolism in cancer: mechanisms and therapeutic opportunities. Nat Metab 2020;2(2):132–141.

Jacobs JJ, Kieboom K, Marino S, DePinho RA, van Lohuizen M. The oncogene and Polycomb-group gene bmi-1 regulates cell proliferation and senescence through the ink4a locus. Nature 1999;397(6715):164-168.

Kallin EM, Cao R, Jothi R, et al. Genome-wide uH2A localization analysis highlights Bmi1-dependent deposition of the mark at repressed genes. PLoS Genet 2009;5(6):e1000506.

Kim AR, Olsen J, England SJ, et al. Bmi-1 regulates extensive erythroid self-renewal. Stem Cell Reports 2015;4(6):995–1003.

Kingsley PD, Malik J, Emerson RL, et al. “Maturational” globin switching in primary primitive erythroid cells. Blood 2006;107(4):1665–1672.

Kobayashi M, Lin Y, Mishra A, et al. Bmi1 maintains the self-renewal property of innate-like B lymphocytes. J Immunol 2020;204(12):3262–3272.

Liu J, Wu M, Lancelot M, et al. BMI1 enables extensive expansion of functional erythroblasts from human peripheral blood mononuclear cells. Mol Ther 2021;29(5):1918–1932.

Liu R, Peng Y, Du W, et al. BMI1 fine-tunes gene repression and activation to safeguard undifferentiated spermatogonia fate. Front Cell Dev Biol 2023;11:1146849.

Lu Z, Huang L, Li Y, et al. Fine-tuning of cholesterol homeostasis controls erythroid differentiation. Adv Sci (Weinh) 2022;9(2):e2102669.

Migliaccio G, Masiello F, Tirelli V, et al. Under HEMA conditions, self-replication of human erythroblasts is limited by autophagic death. Blood Cells Mol Dis 2011;47(3):182–197.

Panzenbock B, Bartunek P, Mapara MY, Zenke M. Growth and differentiation of human stem cell factor/erythropoietin-dependent erythroid progenitor cells in vitro. Blood 1998;92(10):3658–3668.

Park IK, Qian D, Kiel M, et al. Bmi-1 is required for maintenance of adult self-renewing haematopoietic stem cells. Nature 2003;423(6937):302-305.

Qin K, Lan X, Huang P, et al. Molecular basis of polycomb group protein-mediated fetal hemoglobin repression. Blood 2023;141(22):2756–2770.

Rizo A, Olthof S, Han L, Vellenga E, de Haan G, Schuringa JJ. Repression of BMI1 in normal and leukemic CD34+ cells impairs self-renewal and induces apoptosis. Blood 2009;114(8):1498–1505.

Shalev H, Kapelushnik J, Moser A, Knobler H, Tamary H. Hypocholesterolemia in chronic anemias with increased erythropoietic activity. Am J Hematol 2007;82(3):199–202.

Sieweke MH, Allen JE. Beyond stem cells: self-renewal of differentiated macrophages. Science 2013;342(6161):1242974.

Skene P, Henikoff S. An efficient targeted nuclease strategy for high-resolution mapping of DNA binding sites. eLife 2017;6:e21856.

Song LP, Zeng MS, Liao WT, et al. Bmi-1 is a novel molecular marker of nasopharyngeal carcinoma progression and immortalizes primary human nasopharyngeal epithelial cells. Cancer Res 2006;66(12):6225–6232.

Stock JK, Giadrossi S, Casanova M, et al. Ring1-mediated ubiquitination of H2A restrains poised RNA polymerase II at bivalent genes in mouse ES cells. Nat Cell Biol 2007;9(12):1428–1435.

Tan JJ, Wang L, Mo TT, et al. Establishment of immortalized laryngeal epithelial cells transfected with Bmi1. Cell Transplant 2020;29:963689720908198.

van Lohuizen M, Verbeek S, Scheijen B, Wientjens E, van der Gulden H, Berns A. Identification of cooperating oncogenes in E mu-myc transgenic mice by provirus tagging. Cell 1991;65(5):737–752.

von Lindern M, Zauner W, Mellitzer G, et al. The glucocorticoid receptor cooperates with the erythropoietin receptor and c-Kit to enhance and sustain proliferation of erythroid progenitors in vitro. Blood 1999;94(2):550–559.

Wang H, Wang L, Erdjument-Bromage H, et al. Role of histone H2A ubiquitination in Polycomb silencing. Nature 2004;431(7010):873–878.

Wang S, Sun H, Ma J, et al. Target analysis by integration of transcriptome and ChIP-seq data with BETA. Nat Protoc 2013;8(12):2502–2515.

Yan H, Ali A, Blanc L, et al. Comprehensive phenotyping of erythropoiesis in human bone marrow: Evaluation of normal and ineffective erythropoiesis. Am J Hematol 2021;96(9):1064–1076.

Zhang Y, Liu T, Yuan F, et al. The Polycomb protein RING1B enables estrogen-mediated gene expression by promoting enhancer-promoter interaction and R-loop formation. Nucleic Acids Res 2021;49(17):9768–9782.

Zingarriello M, Bardelli C, Sancillo L, et al. Dexamethasone predisposes human erythroblasts toward impaired lipid metabolism and renders their *ex vivo* expansion highly dependent on plasma lipoproteins. Front Physiol 2019;10:281.

