## Supplemental Data for "BMI1 regulates human erythroid self-renewal through both gene repression and gene activation"

#### Supplemental Methods

CUT&RUN

Spearman correlation plots were generated using multiBigwigSummary (bins option) and plotCorrelation (deepTools). Consensus Bmi1 and Ring1b peaks were called using macs2 callpeak (narrowPeak) on merged replicate bam files. Shared/unique peaks were identified using bedtools intersect. Genomic regions were annotated using Homer annotatePeaks.pl. Average enrichment scores were calculated for each mark over promoters (TSS +/- 2kb) using computeMatrix outFileNameMatrix for merged bigwigs (deepTools), and used to generate scatter plots and pearsons correlation values in R. Differentially expressed TSS distance to nearest CpG island was determined using bedtools closest, and parsed by genes with nearest CpG island < 1kb or > 1kb of TSS.

#### Supplemental Figure Legends

**Supplemental Figure 1.** Increased expression of BMI1 increases self-renewal capacity of human erythroid lineage cells.
**(A)** Schema of iSRE establishment and culture initiated from CD34+ cells.
**(B)** Methodology for FACS-based enrichment of self-renewing erythroblasts. iSRE cultures were stained with DAPI (1ug/ml) and sorted using a BD FACSAria II (BD Bioscience) with a 100 um nozzle. Live cells were selected as DAPI$^{-}$, SSC$^{mid\text{-}lo}$ cells, then immature iSRE were selected as FSC$^{hi}$, GFP$^{hi}$ cells.
**(C)** Example of western blot of BMI1 and H4 levels in untransduced self-renewing erythroblasts (UT SRE), empty vector (EV) SREs, and BMI1-transduced (i) SREs.
**(D)** CD34+-derived iSREs expand approximately $10^{12}$-fold. In contrast, untransduced (UT) SREs and empty vector transduced (EV) SREs undergo limited expansion.
**(E)** Normal karyotype of iSREs derived from two independent donors.

**Supplemental Figure 2.** Terminal maturation of human iSREs.
**(A)** Maturation of CD34+-derived iSREs with transition from CD49d+ Band3$^{lo}$ immature erythroblasts to CD49d- Band3+ OrthoE and reticulocytes over 11 days of maturation culture.
**(B)** Example of western blot of BMI1 and control H4 levels at day 0 and day 2 of iSRE maturation.
**(C)** Example of western blots of BMI1 and actin levels following PTC-209-mediated inhibition of BMI1.

**(D)** Dose dependent decrease of BMI1 protein levels following PTC-209 treatment of iSREs. N=6. ***p<0.0003 (2.5uM PTC-209). ****$p<1x10^{-7}$ (12.5uM PTC-209).

**Supplemental Figure 3.** CUT&RUN analysis of BMI1, RING1B, H2AK119Ub, and H3K27me3 in SREs and iSREs.

**(A)** Spearman correlation plots of individual replicates of BMI1, RING1B, H2AK119Ub, and H3K27me3 analyses. One outlier (left-most panel; iSRE-B3) was identified and removed from downstream analyses.

**(B)** Venn diagram of overlap of BMI1 and RING1B peaks in iSREs.

**(C)** Genome annotation of BMI1 peaks show similar distribution in untransduced SREs (left panel) and in BMI1-transduced iSREs (right panel).

**(D)** Pearson correlations of BMI1 peaks at the TSS (± 500bp) are higher with RING1B (upper panels) than with H2AK119Ub marks (lower panels) in untransduced SREs and in iSREs.

**(E)** Pairwise scatterplot analysis of regularized log expression of SRE and iSRE RNA-Seq data revealing high correlation between all samples with SRE and iSRE replicates showing the highest correlation.

**(F)** Volcano plot of genes differentially expressed between SREs and iSREs. N=3 independent samples of each. Upregulated genes include BMI1.

**(G)** BETA analysis reveals statistically significant associations between BMI1-occupied genes and either their positive ($1.8 \times 10^{-23}$) or their negative ($5.5 \times 10^{-11}$) gene expression in iSREs compared to SREs.

**(H)** Heatmap of H2AK119Ub and H3K27me3 in SREs and iSREs at differentially expressed genes sorted by H3K27me3 reveals similar patterns in upregulated and downregulated genes. Color scale represents RPKM of each sample using merged replicates (n=4). Blue line- average peak binding of upregulated genes. Green line- average peak binding of downregulated genes.

**Supplemental Figure 4. Analysis of BMI1 occupancy at CpG islands.**

**(A)** Occupancy of indicated factors and presence of CpG islands in SRE (black) and iSRE (green) at the LIN28b (upper panel), IGF2BP1 (middle panel), and IGF2BP3 (lower panel) loci. Data represent merged replicates (n=4) for all CUT&RUN studies.

**(B)** Expression of LIN28B, IGF2BP1, and IGF2BP3 in SRE and iSRE RNA-seq data sets. There is little expression of LIN28B or IGF2BP1 in these adult derived cells, while IGF2BP3 is significantly downregulated in iSREs with *** adjusted p value $<4e-10^{-10}$.

**(C)** Heatmaps of BMI1, RING1B, H2A119ub, and H3K27me3 occupancy at promoters (2 kb flanking of the TSS) of differentially expressed genes grouped by the presence of CpG island within 1kb of the TSS. Color scale represents RPKM of each sample using merged replicates (n=4). Higher average association of BMI1 and RING1b was seen at promoters with CpG islands in both upregulated genes (dark blue lines) and downregulated genes (green lines) than at promoters of upregulated genes (light blue lines) and downregulated genes (orange lines) genes without CpG islands.

**Supplemental Figure 5.** The BMI1 inhibitor PTC-209 alters cell cycle and increases apoptosis of iSREs.

(**A**) The BMI1 inhibitor PTC-209 decreases the percentage of iSREs in S-phase (BrdU+) in a dose-dependent manner. Representative FACS plots shown.

**(B)** PTC-209 leads to a dose dependent increase in G0 iSREs (Ki67-). Representative FACS plots shown.
**(C)** PTC-209 leads to a dose dependent increase in apoptotic (PI- Annexin5+) iSRE. Representative FACS plots shown.
**(D)** Table of ribosomal protein genes differentially upregulated in iSREs versus SREs. The majority of these ribosomal protein genes (23/33) are also bound by BMI1 at their promoter.

**Supplemental Figure 6.** BMI1 occupies the promoter region of multiple genes regulating key aspects of cholesterol homeostasis.
Occupancy of indicated factors and presence of CpG islands in SRE (black) and iSRE (green) of the indicated loci. Data represent merged replicates (n=4) for all CUT&RUN studies.

### SUPPLEMENTAL FIGURE 1

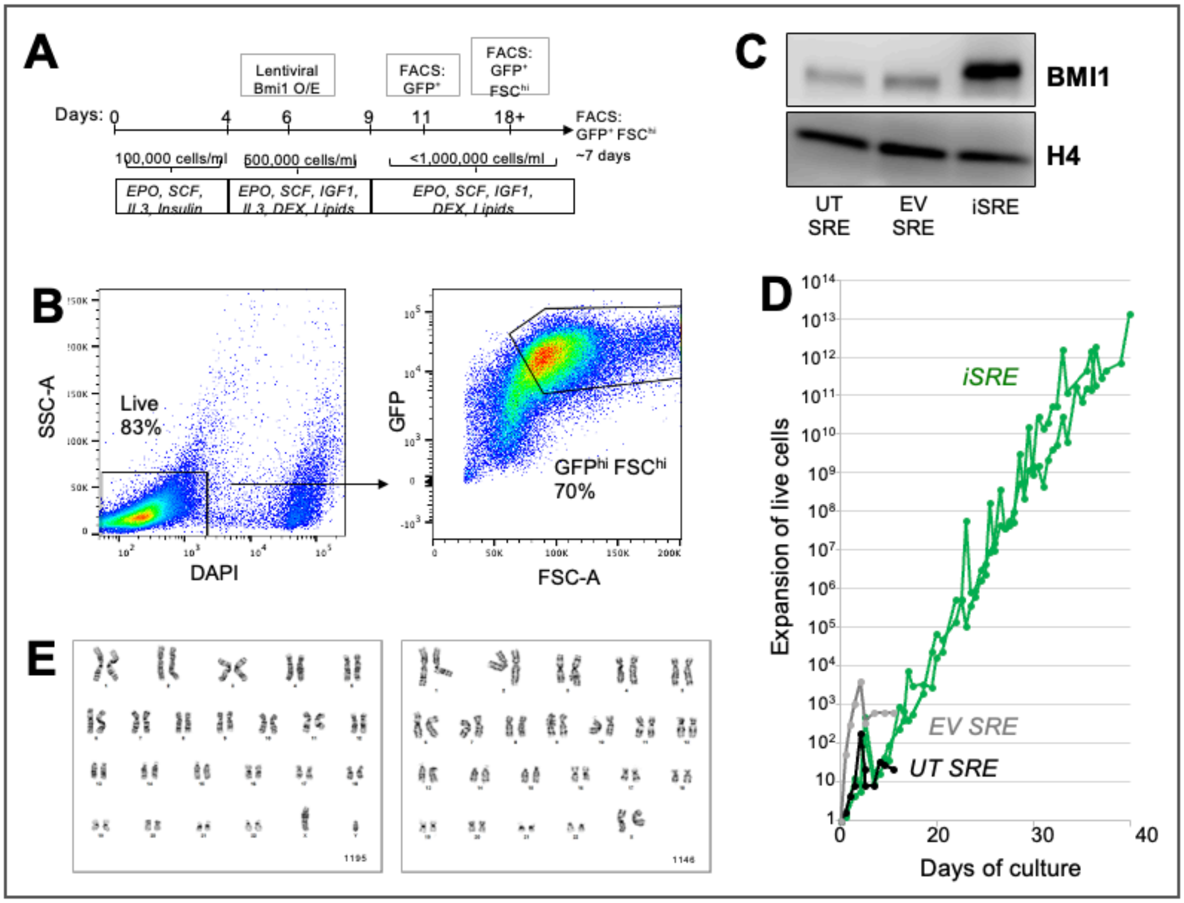

### SUPPLEMENTAL FIGURE 2

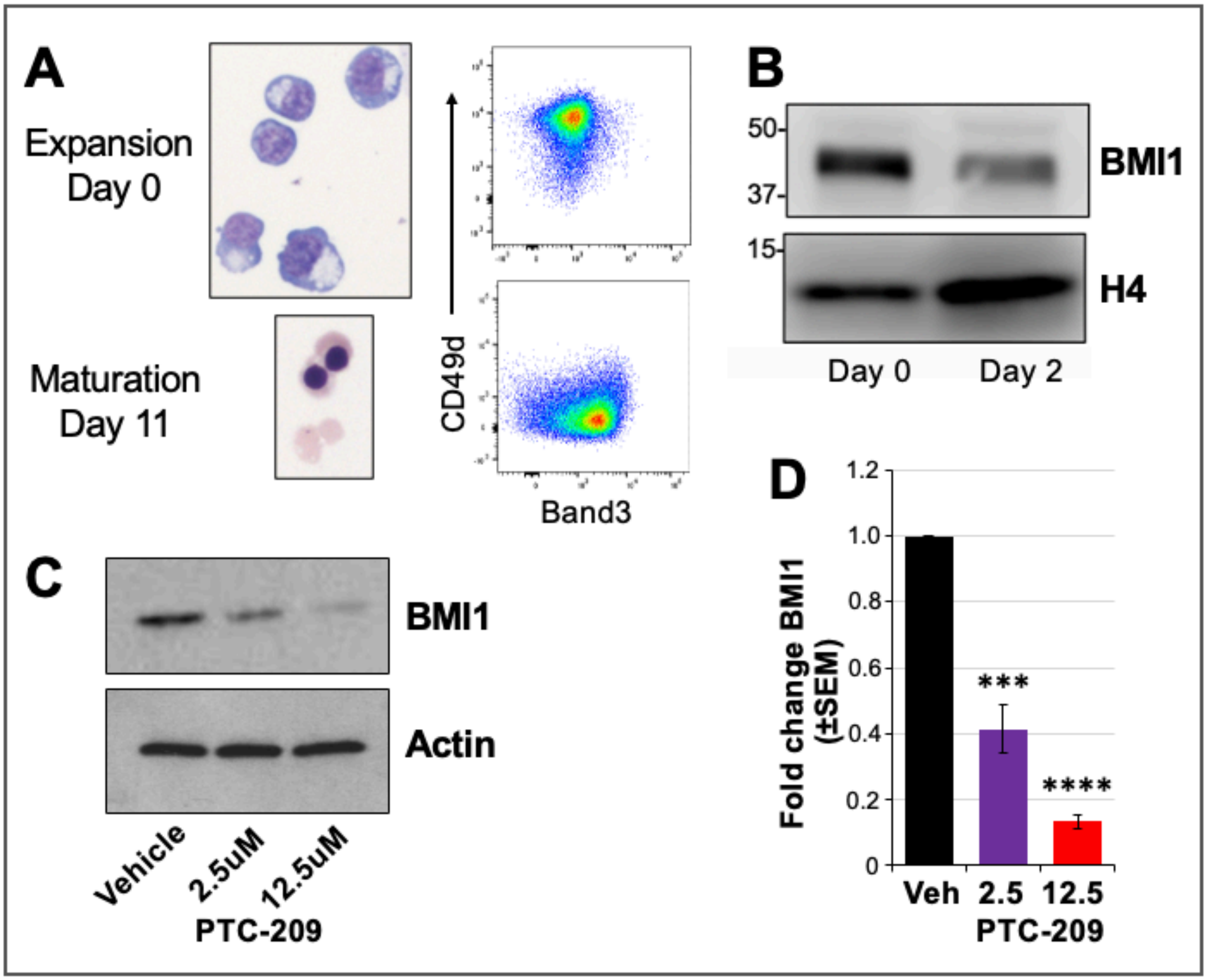

### SUPPLEMENTAL FIGURE 3

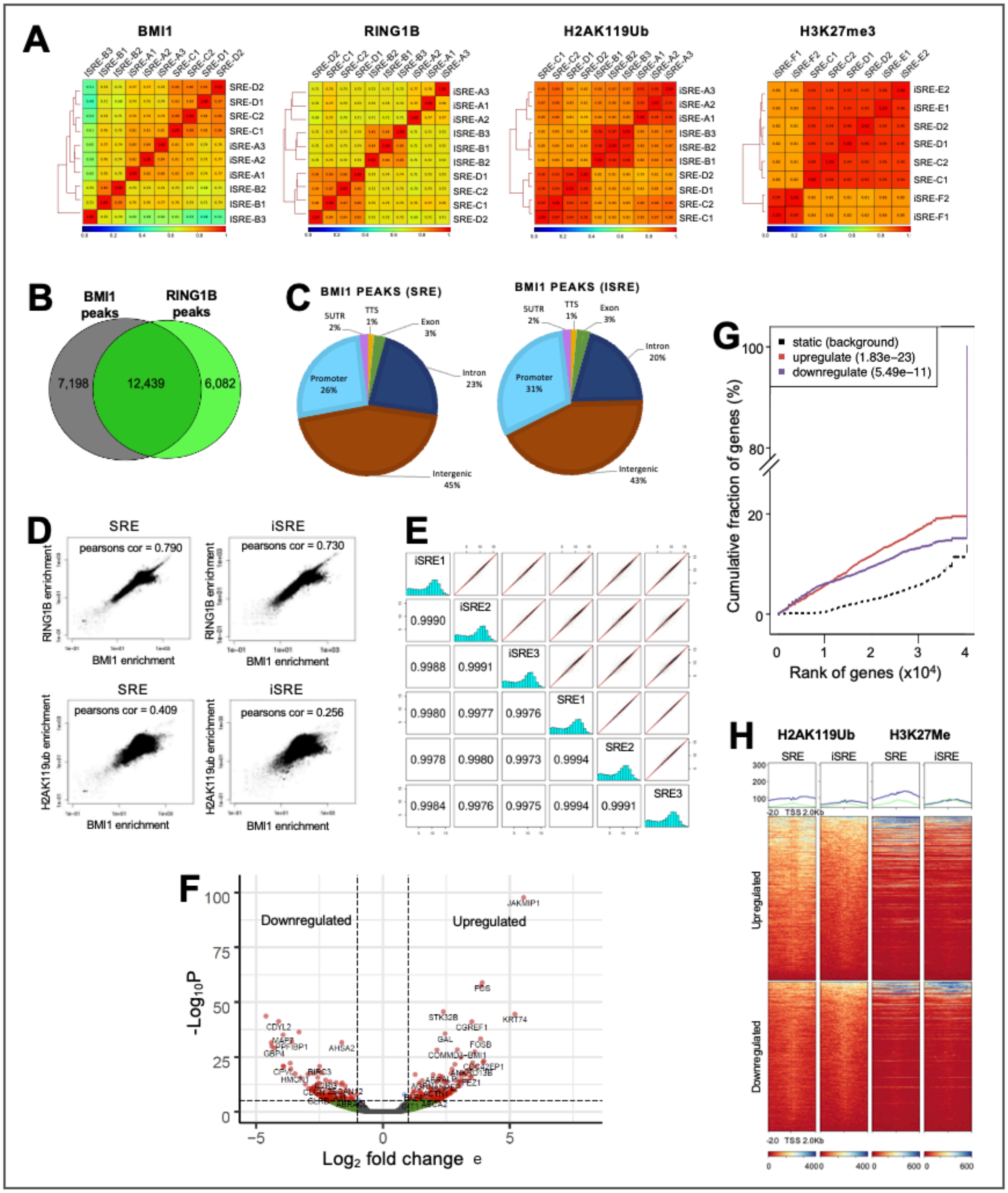

### SUPPLEMENTAL FIGURE 4

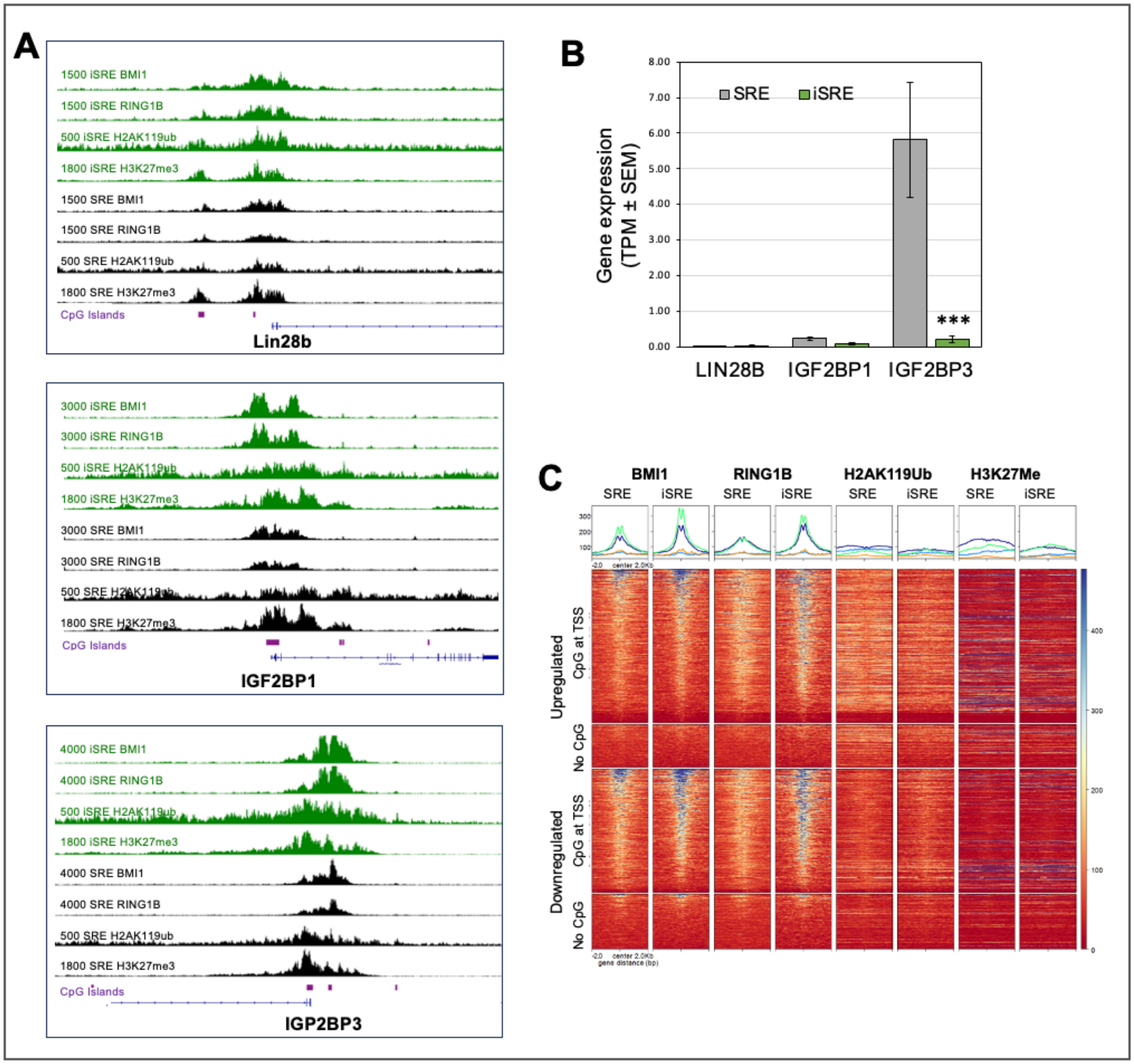

### SUPPLEMENTAL FIGURE 5

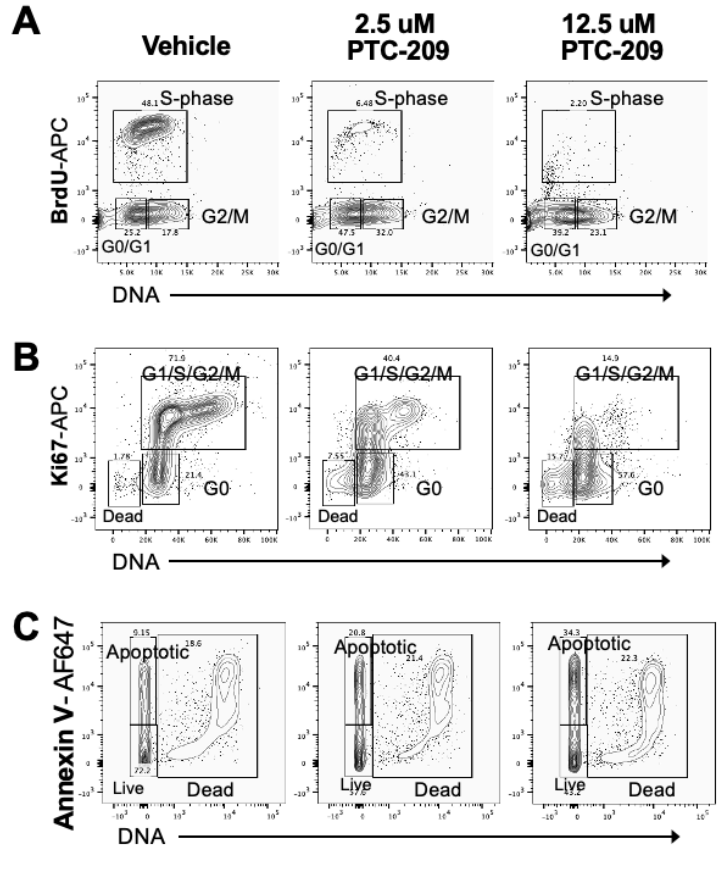


**D**

| | Fold DEG | Adj Pvalue | BMI1 Peak around TSS |
|---|---|---|---|
| Rpl11 | 1.35 | 1.0E-03 | Yes |
| Rpl13a | 1.22 | 4.4E-02 | Yes |
| Rpl15 | 1.31 | 1.4E-03 | |
| Rpl18a | 1.39 | 8.9E-05 | Yes |
| Rpl19 | 1.26 | 1.4E-02 | |
| Rpl22 | 1.23 | 4.0E-02 | Yes |
| Rpl26 | 1.28 | 3.0E-02 | |
| Rpl27 | 1.38 | 1.8E-04 | Yes |
| Rpl27a | 1.41 | 3.9E-03 | Yes |
| Rpl28 | 1.55 | 3.1E-05 | Yes |
| Rpl30 | 1.23 | 4.9E-02 | Yes |
| Rpl32 | 1.35 | 7.0E-04 | Yes |
| Rpl35 | 1.50 | 3.3E-04 | Yes |
| Rpl36 | 1.38 | 2.0E-03 | Yes |
| Rpl7a | 1.32 | 2.8E-03 | |
| Rpl8 | 1.54 | 9.7E-05 | Yes |
| Rplp0 | 1.49 | 2.3E-04 | Yes |
| Rplp1 | 1.27 | 1.6E-03 | Yes |
| Rps10 | 1.36 | 6.3E-04 | |
| Rps11 | 1.41 | 1.3E-03 | Yes |
| Rps14 | 1.27 | 4.5E-02 | Yes |
| Rps15 | 1.35 | 4.0E-04 | |
| Rps15a | 1.40 | 9.6E-04 | |
| Rps16 | 1.36 | 1.1E-06 | Yes |
| Rps17 | 1.45 | 9.0E-04 | Yes |
| Rps19 | 1.35 | 1.6E-03 | Yes |
| Rps2 | 1.55 | 8.5E-07 | |
| Rps23 | 1.23 | 4.0E-02 | Yes |
| Rps26 | 1.52 | 1.6E-02 | Yes |
| Rps27 | 1.31 | 3.2E-03 | |
| Rps28 | 1.36 | 4.6E-06 | |
| Rps5 | 1.37 | 1.0E-03 | Yes |
| Rps9 | 1.32 | 7.6E-03 | Yes |

### SUPPLEMENTAL FIGURE 6

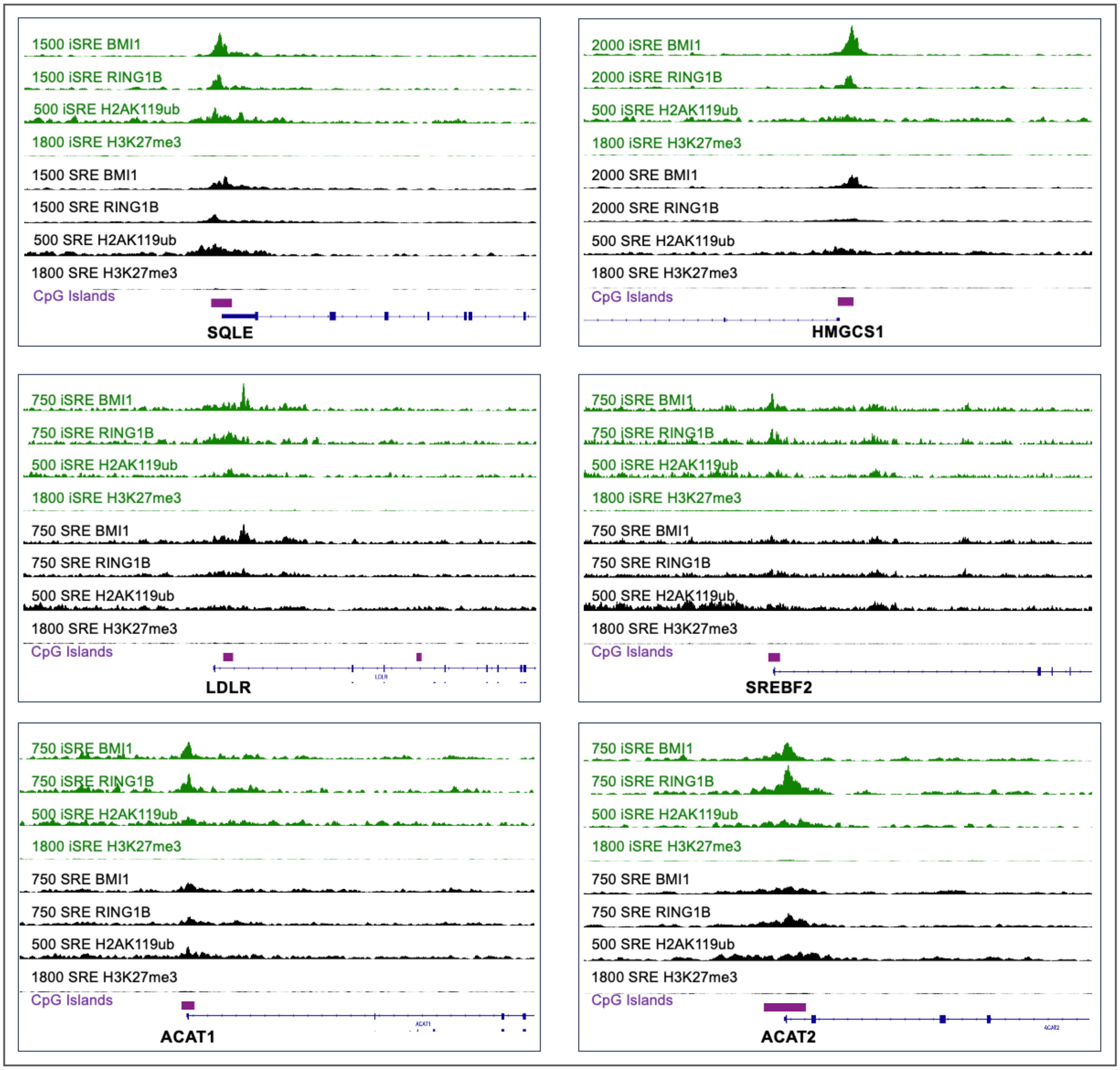

#### Supplemental Tables

**Supplemental Table 1.** Reagents for cell analyses.

| Stain | Figures | Antibody/ Chemical | Fluor | clone | vendor |
|---|---|---|---|---|---|
| SRE/iSRE immunophenotype | 1E; 2B,3B,E; SD2A | CD36 | APC | CB38 | BD Bioscience |
| | | CD235a | PE | HIR2 | Biolegend |
| | | CD71 | APC | OKT9 | Invitrogen |
| | | CD44 | PECF594 | G44-26 | BD Bioscience |
| | | CD49d | PEcf594 | 9F10 | BD Bioscience |
| | | CD14 | PE | MΦP9 | BD Bioscience |
| | | CD11b | PECF594 | ICRF44 | BD Bioscience |
| | | CD34 | APC | 581 | BD Bioscience |
| | | Band 3 | Unstained | BRIC200 | IBGRL |
| | | GPC | Unstained | BRIC10 | IBGRL |
| | | RH-related | Unstained | BRIC69 | IBGRL |
| | | CD235a | Unstained | BRIC256 | IBGRL |
| | | CD238 Kell | Unstained | BRIC203 | IBGRL |
| | | 2° Antimouse | PE | | ThermoFisher |
| | | 2° Antimouse | AF647 | | ThermoFisher |
| | | DAPI | | | Sigma |
| | | DRAQ5 | | | ThermoFisher |
| Cell cycle Apoptosis | 7C,D,E; SD5A,B,C | anti-BrDU (Flow kit) | FITC | | BD Bioscience |
| | | Ki67 | APC | Ki-67 | Biolegend |
| | | Annexin V | AF647 | | ThermoFisher |
| | | 7AAD | | | ThermoFisher |
| | | Propidium Iodide | | | ThermoFisher |
| Lipid Analysis | 8F,G | CD235a | BV421 | GA-R2 (HIR2) | BD Bioscience |
| | | CD105 | PE | 266 | BD Bioscience |
| | | CD45RA | PEcy7 | HI100 | BD Bioscience |
| | | CD41a | PEcy7 | HIP8 | BD Bioscience |
| | | CD123 | PEcy7 | 7G3 | BD/SC |
| | | CD14 | PEcy7 | M5E2 | BD Bioscience |
| | | CD16 | PEcy7 | eBioCB16 (CB16) | eBioscience |
| | | CD71 | APC-H7 | M-A712 | BD Bioscience |
| | | Cholera Toxin Beta (CB-T) | FITC | | Sigma-Aldrich |
| | | Filipin | | | Sigma |

**Supplemental Table 2.** Taqman probes for qPCR.

| Gene | Taqman. Probe |
|---|---|
| BMI1 | Hs00180411_m1 |
| CDKN1A | hs00355782_m1 |
| CDKN1B | hs00153277_m1 |
| CDKN2A-INK4A | hs02902543_mH |
| CDKN2A-ARF | hs99999189_m1 |
| CDKN2B | hs00793225_m1 |
| CDKN2C | hs00176227_m1 |
| CDKN2D | hs00176481_m1 |
| HBB | Hs00758889_m1 |
| HBG1/HBG2 | Hs00361131_g1 |
| HBA1/HBA2 | Hs00361191_g1 |
| HBE1 | HS00362216_m1 |
| HBZ | HS00923579_m1 |
| 18s | Hs99999901_s1 |
| HMGCR | Hs00168352_m1 |
| HMGCS1 | Hs00940429_m1 |
